# Heterogeneous profiles of coupled sleep oscillations in human hippocampus

**DOI:** 10.1101/589978

**Authors:** Roy Cox, Theodor Rüber, Bernhard P Staresina, Juergen Fell

## Abstract

Cross-frequency coupling of sleep oscillations is thought to mediate memory consolidation. While the hippocampus is deemed central to this process, detailed knowledge of which oscillatory rhythms interact in the sleeping human hippocampus is lacking. Combining intracranial hippocampal and non-invasive electroencephalography from twelve neurosurgical patients, we characterized spectral power and coupling during non-rapid eye movement (NREM) and rapid eye movement (REM) sleep. Hippocampal coupling was extensive, with the majority of channels expressing spectral interactions. NREM consistently showed delta–ripple coupling, but ripples were also modulated by slow oscillations (SOs) and sleep spindles. SO–delta and SO–theta coupling, as well as interactions between delta/theta and spindle/beta frequencies also occurred. During REM, limited interactions between delta/theta and beta frequencies emerged. Moreover, oscillatory organization differed substantially between i) hippocampus and scalp, ii) sites along the anterior-posterior hippocampal axis, and iii) individuals. Overall, these results extend and refine our understanding of hippocampal sleep oscillations.

## 1 Introduction

The hippocampus and wider medial temporal lobe system play a crucial role in episodic memory, the ability to recall past events and their individual components (Squire et al., 2004; Tulving and Markowitsch, 1998). Besides its role in memory formation and retrieval, the hippocampus (HPC) is also a key player in systems consolidation, whereby initially labile memories are gradually transferred from HPC to neocortical sites for permanent storage (Buzsáki, 1996; Frankland and Bontempi, 2005; Marr, 1971; Zola-Morgan and Squire, 1990). Intriguingly, consolidation depends heavily on sleep (Ellenbogen et al., 2006; Jenkins and Dallenbach, 1924; Walker et al., 2002), with greatest importance usually ascribed to non-rapid eye movement (NREM) sleep, and a relatively poorly understood role for rapid eye movement (REM) sleep (Rasch and Born, 2013; Stickgold and Walker, 2013).

NREM-dependent consolidation is thought to rely on the unique electrophysiological signatures of this brain state. These include 0.5–1 Hz neocortical slow oscillations (SOs), 12.5–16 Hz thalamocortical fast sleep spindles, and 80–100 Hz hippocampal ripples (frequency ranges in humans), all of which have been associated with human memory reactivation and consolidation processes (Axmacher et al., 2008; Cairney et al., 2018; R. Cox et al., 2014; Huber et al., 2004; Schönauer et al., 2017; Zhang et al., 2018). Although SOs and spindles are most prominent in neocortex, they are also present in human HPC (Andrillon et al., 2011; Frauscher et al., 2015; Nir et al., 2011; Staresina et al., 2015), consistent with their postulated role in HPC-cortical memory transfer. Interestingly, SOs, spindles, and ripples are temporally organized, with faster oscillations preferentially expressed at a particular phase of the slower rhythm. In general terms, such cross-frequency phase-amplitude coupling (PAC) is thought to enable brain communication across multiple spatiotemporal scales (Aru et al., 2015; Canolty and Knight, 2010). During sleep, PAC occurs both within and across brain areas (Cox et al., 2018; R. Cox et al., 2014; Klinzing et al., 2016; Mak-McCully et al., 2017; Mölle et al., 2011), including in human (para)hippocampus (Clemens et al., 2011, 2007; Staresina et al., 2015). Remarkably, animal findings have shown that stronger SO–spindle–ripple coupling enhances plasticity and memory retention (Latchoumane et al., 2017; Maingret et al., 2016; Niethard et al., 2018), suggesting that these nested oscillations are required for effective information transfer and storage (Born et al., 2006).

In addition to these “classical” forms of NREM PAC among SOs, spindles, and ripples, SOs affect the expression of 1–4 Hz delta activity (Steriade et al., 1993), which is physiologically distinct from SOs (Achermann and Borbély, 1997; Amzica and Steriade, 1998; Benoit et al., 2000). Whether SO–delta coupling also exists in human HPC, and whether HPC SO and delta activity coordinate faster activity in a similar fashion, is presently unknown. Moreover, HPC ripples often occur in conjunction with ~4 Hz sharp waves (Buzsáki, 1986), which could be expressed as delta–ripple PAC (Oliva et al., 2018; Staresina et al., 2015). Finally, neocortical theta (~6 Hz) activity is also organized by the SO phase (Gonzalez et al., 2018), and this coupling has been related to memory consolidation (Schreiner et al., 2018). Whether analogous SO–theta coupling exists in human HPC is unknown.

Contrary to the relatively organized structure of NREM oscillations, REM sleep exhibits an overall cortical “desynchronization” (i.e., fast, low-amplitude activity), although isolated REM SOs have been described in rodent cortex (Funk et al., 2016). Similarly, human HPC expresses both fast beta/gamma activity (Uchida et al., 2001) and relatively slow activity during REM sleep. Yet, whether this slower component entails SO/delta (Bódizs et al., 2001) or theta (Cantero et al., 2003) activity is still contentious. Moreover, with the exception of delta–gamma coupling in parahippocampal areas (Clemens et al., 2009), the extent of interacting HPC oscillations in human REM sleep has not been explored.

Importantly, the HPC consists of anatomically and functionally distinct domains, with anterior and posterior regions showing differential relations with human memory performance (Bonnici et al., 2012; Dandolo and Schwabe, 2018; Ludowig et al., 2008; Poppenk and Moscovitch, 2011). Given that sleep oscillations are often spatially restricted (Cox et al., 2018; R. Cox et al., 2014; Nir et al., 2011; Vyazovskiy et al., 2011), it is an open question whether these rhythms and their interactions vary along the longitudinal axis of the human HPC, possibly in relation to functional specialization. Another relevant factor concerns between-subject variability, with various aspects of scalp-recorded sleep oscillations showing large and reproducible individual differences (Cox et al., 2018, 2017; De Gennaro et al., 2005; Massimini et al., 2004; Rusterholz et al., 2018). Whether such variability extends to human HPC is unknown.

In sum, while electrophysiological HPC activity during sleep is deemed pivotal for memory processing, a comprehensive characterization of human HPC sleep oscillations and their interactions has not been performed. The wealth of fundamental animal knowledge notwithstanding (Mizuseki and Miyawaki, 2017; Timofeev, 2011; Todorova and Zugaro, 2018), non-negligible species differences in the expression and frequency of sleep oscillations make it unclear whether theoretical accounts on the role of the sleeping HPC are consistent with human data. Taking advantage of the unique opportunity offered by neurosurgical monitoring in 12 epilepsy patients, we explored oscillatory activity along the longitudinal HPC axis during light NREM (N2), deep NREM (N3), and REM sleep. For comparison purposes, we also included scalp activity. We assessed spectral power and PAC across an extensive (0.5–200 Hz) frequency range, employing data-driven permutation-based approaches to characterize and contrast oscillatory activity in various ways. Of note, our PAC approach considered continuous data rather than focusing on a priori defined brain rhythms or graphoelements, thus allowing detection of coupling effects outside the common SO-spindle-ripple framework. Our results indicate extensive PAC in human HPC, with the majority of channels expressing clear spectral interactions, but also a remarkable heterogeneity between HPC and scalp, HPC sites, and individuals.

## 2 Materials and Methods

### 2.1 Participants

We analyzed archival electrophysiological sleep data in a sample of 12 (6 male) patients suffering from pharmaco-resistant epilepsy (age: 36.5 ± 14.5 yrs, range: 22–62). This sample overlaps with ones presented previously (Staresina et al., 2015; Wagner et al., 2010). Patients had been epileptic for 23.4 ± 12.0 yrs (range: 10–49) and were receiving anticonvulsive medication at the moment of recording. All patients gave informed consent, the study was conducted according to the Declaration of Helsinki, and was approved by the ethics committee of the Medical Faculty of the University of Bonn.

### 2.2 Data acquisition

Electrophysiological monitoring was performed with a combination of depth and subdural strip/grid electrodes placed according to clinical criteria. HPC depth electrodes (AD-Tech, Racine, WI, USA) containing 8–10 cylindrical platinum-iridium contacts (length: 1.6 mm; diameter: 1.3 mm diameter; center-to-center inter-contact distance: 4.5 mm) were stereotactically implanted. Implantations were done either bilaterally (n=9) or unilaterally (n=3), and either along the longitudinal HPC axis via the occipital lobe (n=11) or along a medial-lateral axis via temporal cortex (n=1).

Pre- and post-implantation 3D T1-weighted magnetic resonance image (MRI) scans were used to determine electrode locations. Pre-operative T1 (resolution = 0.8×0.8×0.8 mm^3^, TR = 1,660 ms, TE = 2.54 ms, flip angle = 9°) was acquired using a 3.0 Tesla Magnetom Trio (Siemens Healthineers, Erlangen, Germany) with a 32-channel-coil. Post-operative T1 (resolution = 1×1×1 mm^3^, TR = 11.09 ms, TE = 5.02 ms, flip angle = 8°) was conducted using an Achieva 3.0 Tesla Tx system (Philips Healthcare, Best, The Netherlands). Preprocessing and analyses of T1 volumes was done using FMRIB’s Software Library 5.0 (FSL) (Jenkinson et al., 2012). Brain extractions (Smith, 2002) were performed and followed by a bias-field correction (Zhang et al., 2001). Post-operative volumes were linearly registered to the pre-operative volumes. Pre-operative volumes were normalized to the Montreal Neurological Institute template (MNI 152, 1 mm resolution) by means of a non-linear registration (Jenkinson and Smith, 2001). The resulting normalization matrices were then applied to the registered post-operative volumes to normalize them to the MNI template. This way, the normalization process was not affected by electrode image artifacts. Anatomical labels of the electrodes were determined by an experienced physician (TR) based on the subject-specific co-registered T1 volumes. MNI coordinates were extracted from the normalized volumes.

We included a total of 61 HPC contacts (5.1 ± 2.3 per patient, range: 1–8). Only electrodes from the non-pathological side were considered (6 left), and electrode contacts were included only when they could be localized to HPC gray matter, the HPC gray/white matter border, or the border between HPC gray matter and the lateral ventricle. A manual subdivision was made into anterior (total: 23, range: 0–4), middle (22, 0–3), and posterior (16, 0–3) thirds of the HPC. Note that not every patient had contacts inside each hippocampal subdivision (see Supp. Table 1 for an overview of anatomical labels and MNI coordinates for all electrodes). Anterior (73.7 ± 2.6 mm) and posterior (75.8 ± 3.6 mm) contacts were equally distant to the ipsilateral mastoid process (t(7)=-2.0, P=0.09), with average differences of only 2.1 ± 3.0 mm (range: −2.6 to 7.9) as determined from MNI coordinates. For sleep recordings, additional signals were recorded from the scalp (Cz, C3, C4, Oz, A1, A2; plus T5 and T6 in 10 patients), the outer canthi of the eyes for electrooculography (EOG), and chin for electromyography (EMG). All signals were sampled at 1 kHz (Stellate GmbH, Munich, Germany) with hardware low- and high-pass filters at 0.01 and 300 Hz respectively, using an average-mastoid reference.

### 2.3 Sleep scoring

Offline sleep scoring was done in 20 s epochs based on scalp EEG, EOG, and EMG signals in accordance with Rechtschaffen and Kales criteria (Rechtschaffen and Kales, 1968). Stages S3 and S4 were combined into a single N3 stage following the more recent criteria of the American Academy of Sleep Medicine (Silber et al., 2007).

### 2.4 Preprocessing and artifact rejection

All data processing and analysis was performed in Matlab (the Mathworks, Natick, MA), using custom routines and EEGLAB functionality (Delorme and Makeig, 2004). For each patient, we included all epochs scored as sleep or wake and selected channels of interest (including channels not reported in the present paper). Data were filtered with a repeated notch filter (50 Hz and harmonics up to 300 Hz), and a 0.3 Hz high-pass. In a first round of artifact rejection, we visually identified and marked time segments during which any channel displayed obvious artifacts (e.g., excessive amplitudes, amplifier saturation). Next, remaining time segments were used to determine channel-specific thresholds for detection of epileptogenic artifacts. Specifically, each time point was converted to a z-score based on the mean and SD of i) the signal gradient (amplitude difference between adjacent time points), and ii) the signal amplitude after applying a 250 Hz high-pass filter. Artifacts were marked from 0.5 s before to 0.5 s after contiguous time points where the z-score of either of these measures exceeded a critical value. All records were visually reviewed to confirm that threshold settings resulted in appropriate artifact detection. We found that a z threshold of 6 offered an optimal balance between detection of epileptogenic spikes and unwarranted rejection of clean data. Next, we included only continuous data segments that were free of either type of artifact across all included channels for a minimum of 3 s. That is, if a single channel displayed an artifact, that data segment would not be included for any of the channels, further reducing the likelihood that data is contaminated by pathological activity. Data within 0.5 s from non-adjacent epochs was also excluded. All told, this approach removed 54.6 ± 16.6% of the data, leaving a total of 86.7 ± 41.3 (N2), 26.3 ± 25.5 (N3), and 55.3 ± 35.8 min (REM) of clean data. Remaining segments (“trials”) were split if they spanned multiple sleep/wake stages and if the resulting new trials also exceeded a duration of 3 s. Each trial was assigned a stage label based on its midpoint. Average number and duration of included trials was 763 ± 383 and 8.3 ± 7.4 s (N2), 241 ± 210 and 6.6 ± 3.7 s (N3), and 272 ± 172 and 16.1 ± 12.7 s (REM).

### 2.5 Spectral analysis

For each trial and channel, we estimated power spectral density using Welch’s method with 3 s windows and 80% overlap (0.244 Hz resolution). Mean stage spectra were determined with a weighted average approach using trial durations as weights. Next, we removed the spectra’s 1/f component to better emphasize narrowband spectral peaks. To this end, we first interpolated the notch-filtered region (50, 100, 150, and 200 Hz, ± 5 Hz) of each spectrum (Modified Akima cubic Hermite algorithm). Then, we took the N2 spectrum of Cz and the channel-averaged N2 spectrum of the HPC, and fit each according to *af*^*b*^ using log-log least squares regression (Miller et al., 2009; Vijayan et al., 2017). Fitting range was restricted to the 4–175 Hz range to avoid the often observed flattening of the spectrum below ~4 Hz (see Fig. 2AB insets) and the ~200 Hz interpolated data. Then, for each channel and stage, the N2 model fit was subtracted from the observed spectrum. We applied the N2 fit to all stages rather than using stage-specific fits to enable direct stage comparisons. Similarly, for HPC channels we subtracted the channel-averaged spectrum from each channel to allow channel comparisons. Adjusted spectra were resampled to log space and smoothed three times with a moving average window of length 5.

### 2.6 Time-frequency decomposition

In order to assess sleep oscillatory dynamics across several orders of magnitude, we decomposed the multichannel data with a family of complex Morlet wavelets. Each trial was extended with 5 s on either side to minimize edge artifacts. Wavelets were defined in terms of desired temporal resolution according to:

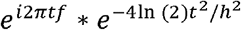

where *i* is the imaginary operator, *t* is time in seconds, *f* is frequency (50 logarithmically spaced frequencies between 0.5 and 200 Hz), *ln* is the natural logarithm, and *h* is temporal resolution (full-width at half-maximum; FWHM) in seconds (Cohen, 2019). We set *h* to be logarithmically spaced between 3 s (at 0.5 Hz) and 0.025 s (at 200 Hz), resulting in FWHM spectral resolutions of 0.3 and 35 Hz, respectively. Trial padding was trimmed from the convolution result and data were downsampled by a factor four to reduce the amount of data. We assessed PAC using a surrogate approach (see below). To make surrogate distributions independent of variable numbers and durations of trials, we first concatenated the convolution result of all trials of a given sleep stage, and then segmented them into 1 min fragments (discarding the final, incomplete segment). Raw PAC and surrogate distributions were then determined separately for each 1 min segment. Segment length was chosen to obtain stable PAC estimates that are not overly affected by cycle-by-cycle variations in coupling strength and/or phase.

### 2.7 Cross-frequency coupling

PAC was determined between all pairs of modulating frequency *f1* and modulated frequency *f2*, where *f2*>2\**f1*. We employed an adaptation of the mean vector length method (Canolty et al., 2006) that adjusts for possible bias stemming from non-sinusoidal shapes of *f1* (van Driel et al., 2015). Specifically, complex-valued debiased phase-amplitude coupling (dPAC) was calculated as:

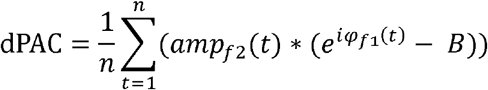

where *i* is the imaginary operator, *t* is time, *amp*_*f2*_(*t*) is the magnitude of the convolution result, or amplitude, of *f2*, *φ*_*f1*_(*t*) is the phase of *f1*, and *B* is the mean phase bias:

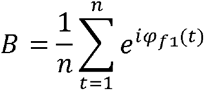

Raw coupling strength (i.e., the degree to which the *f2* amplitude is non-uniformly distributed over *f1* phases) was defined as the magnitude (i.e., length) of the mean complex vector. For every 1 min segment, channel, and frequency pair, we constructed a surrogate distribution of coupling strengths by repeatedly (n = 100) time shifting the *f1* phase time series with respect to the *f2* amplitude time series by a random amount between 1 and 59 s, and recalculating the mean vector length for each iteration. Note that time shifting is a more conservative approach than fully scrambling time series, which may result in spurious effects (Scheffer-Teixeira and Tort, 2016). We then z-scored the observed coupling strength with respect to this null distribution of coupling strength values. Thus, the z-scored measure (dPAC_Z_) indicates how far, in terms of standard deviations, the observed coupling estimate is removed from the average coupling estimate under the null hypothesis of no coupling. All reported analyses of coupling strength are based on dPAC_Z_, but for simplicity we refer to this measure as PAC. Coupling phase (i.e., the *f1* phase where *f2* amplitude is maximal) was defined as the phase angle of the mean complex vector.

### 2.8 Statistics

Statistical analyses were performed at three hierarchical levels of inference: the channel, the subject, and the group level. Specifically, power was analyzed at the subject and group (but not channel) levels, whereas PAC was analyzed at the channel and group (but not subject) levels. We used cluster-based permutation statistics (Maris and Oostenveld, 2007) to determine the presence of PAC, and to compare power and PAC between sleep stages or brain regions. Briefly, a t-test was performed at each frequency bin (power) or frequency-frequency bin (PAC), and bins meeting a threshold (P=0.1) were allowed to form clusters based on spectral adjacency (one-dimensional for power, two-dimensional for PAC). T-values within each cluster were summed as a measure of effect size (cluster statistic). Next, condition labels were repeatedly shuffled and the largest cluster statistic at each iteration was stored. Number of iterations depended on the number of possible permutations: If the number of possible combinations was over 1,000, a random shuffle was performed for each of 1,000 iterations (i.e., Monte Carlo sampling), otherwise each unique combination was sampled exactly once (i.e., permutation sampling). Observed cluster statistics were then compared to the surrogate distribution of cluster statistics. Clusters were considered significant at P<0.05 for one-sided tests (presence of coupling), and at P<0.025 for two-sided tests (condition comparisons).

At the channel level (PAC only), the presence of coupling (e.g., Fig. 3A) was determined by comparing the distribution of dPAC_Z_ values across segments to zero (paired t-test). Channel and stage differences (e.g., Fig. 3CG) were assessed using paired and unpaired t-tests, respectively. At the subject level (power only), trial-averaged spectra were compared between sleep stages across channels (e.g., Fig. 2DG, paired t-test). At the group level, power/dPAC_Z_ values were first averaged across trials/segments, and, for HPC, relevant channels (i.e., anterior, middle, posterior, or all channels). The distribution of these values was compared to zero (e.g., Fig. 5A; paired t-tests), and contrasted between brain regions and stages (e.g., Fig. 2A, 4BC; paired t-tests).

Since our objective was to offer a comprehensive characterization of the oscillatory structure of human HPC, this study should be viewed as a hypothesis-generating (or exploratory), rather than a hypothesis-testing (or confirmatory) study. Consequently, while we employed rigorous permutation-based approaches for every analysis, we did not further adjust for multiple testing (e.g., no correction across all panels of Fig. 4AB).

### 2.9 Density-based clustering

To obtain a qualitative picture of how many individuals showed PAC for each frequency pair we employed the *density-based spatial clustering of applications with noise* (DBSCAN) algorithm (Ester et al., 1996), an unsupervised clustering method that aggregates nearby points without requiring an *a priori* specification of the number of clusters. In detail, for every individual, sleep stage, and channel, we extracted the frequency-frequency coordinates where dPAC_Z_ was at a local maximum. Note that local maxima were generally of larger magnitude for NREM than REM, allowing for the inclusion of weaker effects for REM. Local maxima carried forward to the clustering approach were selected based on both an adaptive and a fixed threshold approach. For the adaptive approach, maxima were rank ordered and thresholded according to: *prop*_*inc*_ = *c* × *N*_*chan*_, where *prop*_*inc*_ is the proportion of local maxima that is included, c is a thresholding coefficient between 0 and 1, and *N*_*chan*_ is the number of channels within a region. We set c to 0.3, but obtained similar results with values of 0.2 and 0.4. *N*_*chan*_ always equaled 1 at the scalp, but ranged from 1 to 8 for HPC. Hence, the adjustment for *N*_*chan*_ ensures we included approximately similar numbers of dPAC_Z_ values (and corresponding frequency coordinates) for each subject, but also that each HPC channel was allowed to contribute its strongest effects. For the fixed approach, only local maxima with a dPAC_Z_ score above 2 were included. Importantly, to allow large local maxima that were only present on a single channel to survive the clustering approach, frequency-frequency points were duplicated according to their dPAC_Z_ value. Specifically, above-threshold dPAC_Z_ scores were divided into 5 equally sized bins (ranging from the threshold value to the largest dPAC_Z_ score), and each frequency-frequency point was multiplied n times according to the bin number containing its dPAC_Z_ score. Then, for each individual, sleep stage, and region, resulting frequency coordinates were submitted to the DBSCAN algorithm. Frequency coordinates were assigned to the same cluster if they were a maximum distance of 1.5 frequency-frequency bins away, and clusters were required to contain a minimum of 2 points (similar results with minimum cluster sizes of 3 and 4). Frequency coordinates not meeting these criteria were labeled as noise and discarded. We extracted each cluster’s center of gravity as the dPAC_Z_-weighted average of the frequency coordinates of that cluster’s members, and assigned it to the closest discrete frequency-frequency bin. Examples of single-subject cluster assignment are presented in Supp. Fig. 4. A binary frequency-frequency image was generated for every individual, sleep stage, and region, with ones at cluster centers and zeros elsewhere. Binary images were smoothed with a 2D Gaussian filter of standard deviation 0.5, rounded upwards so as to consist of only zeros and ones, and summed across subjects to generate Fig. 6.

### 2.10 Data and code availability

Data are not publicly available due to privacy concerns related to clinical data, but data and accompanying analysis code are available from the corresponding or senior author upon obtaining ethical approval.

## 3 Results

We examined overnight scalp and invasive hippocampal electroencephalography (EEG) in a sample of 12 epilepsy patients. A total of 61 HPC contacts were included (5.1 ± 2.3 per patient, range: 1–8). Only HPC contacts from the non-pathological hemisphere were used, as evidenced by clinical monitoring. HPC contacts were classified as anterior, middle, or posterior HPC based on individual anatomy (see Supp. Table 1 for all electrode locations). Polysomnography-based sleep architecture is shown in Table 1.

**Table 1.** Sleep architecture. WASO: wake after sleep onset.

|  | mean | SD |
| --- | --- | --- |
| N1(%) | 29.8 | 21.8 |
| N2 (%) | 40.5 | 13.7 |
| N3 (%) | 13.2 | 9.0 |
| REM (%) | 16.5 | 8.0 |
| N1 (min) | 171.4 | 175.0 |
| N2 (min) | 203.9 | 74.0 |
| N3 (min) | 64.1 | 45.7 |
| REM (min) | 81.1 | 41.0 |
| total sleep (min) | 520.5 | 113.5 |
| WASO (min) | 82.9 | 77.2 |
| sleep efficiency (%) | 87.1 | 11.7 |

### 3.1 Raw traces and spectrograms

Fig. 1 shows 10 s of concurrent scalp (Cz) and HPC traces and spectrograms for a single participant during N2, N3, and REM sleep. Scalp observations (top) were in line with typical sleep, with prominent NREM fast spindles (most clearly in N2), N3 SO/delta activity, and mixed frequency activity during REM. While HPC followed this general trend (bottom), HPC and scalp activities were also distinct. Bursts of NREM spindle activity were often synchronous (pairs of white arrows), but could be stronger either at the scalp (yellow) or in HPC (gray). At other times, spindle activity appeared exclusively in one site (magenta). Intriguingly, we observed instances of very prominent HPC spindles during REM sleep that were wholly absent at the scalp (green). Moreover, several other spectral components could be discerned, including in the delta range (orange), as well as the theta, beta, and ripple ranges (examples in Supp. Fig. 1), depending on patient, channel and sleep stage. These observations illustrate the diverse constellation of oscillatory rhythms that contribute to the observed signal.

**Figure 1.**
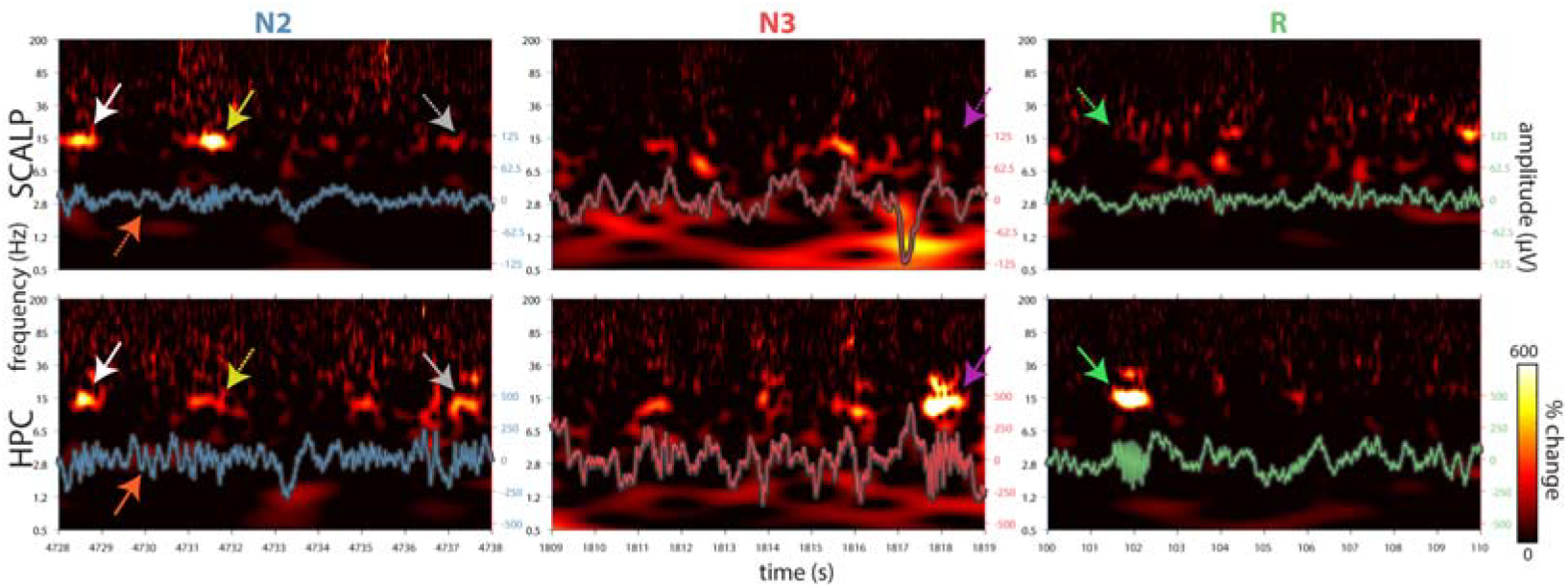
Examples of scalp and hippocampal sleep activity. Shown are 10 s segments of artifact-free data from a single patient (p4). Note the different amplitudes of scalp and hippocampal traces. Spectrograms show percent amplitude change relative to mean across all artifact-free N2, N3 and REM sleep. Arrows illustrate phenomena described in the main text (solid: stronger effect; dashed: weaker effect). Hippocampal channel located in posterior hippocampus (TL08 in Fig 4AC).

### 3.2 Spectral profiles

Considering a total of 86.7 ± 41.3 (N2), 26.3 ± 25.5 (N3), and 55.3 ± 35.8 min (REM) of artifact-free data, power spectra were determined for each channel and sleep stage, adjusting for 1/f power scaling with a slope-fitting procedure to enhance the visibility of narrowband peaks. Averaged across subjects, the scalp spectrum showed greatest SO/delta (0.5–4 Hz) power in N3, prominent fast spindle (12.5–16 Hz) and theta (4–8 Hz) peaks in both NREM stages, and theta and beta (~25 Hz) peaks in REM (Fig. 2A), consistent with typical sleep (Cox et al., 2017; De Gennaro et al., 2005). The HPC spectrum also displayed a strong fast spindle peak and additional delta/theta peaks during NREM (Fig. 2B). Most prominently, HPC showed a strong gamma (40–100 Hz) peak in all sleep stages, matching earlier observations in parahippocampal cortex (Uchida et al., 2001) and consistent with the frequency range of human ripples (Axmacher et al., 2008; Staresina et al., 2015). Results of sleep stage comparisons are indicated at the top of each plot. Direct comparisons between scalp and HPC confirmed the specificity of HPC gamma while showing greater <20 Hz power at the scalp (Supp. Fig. 2).

**Figure 2.**
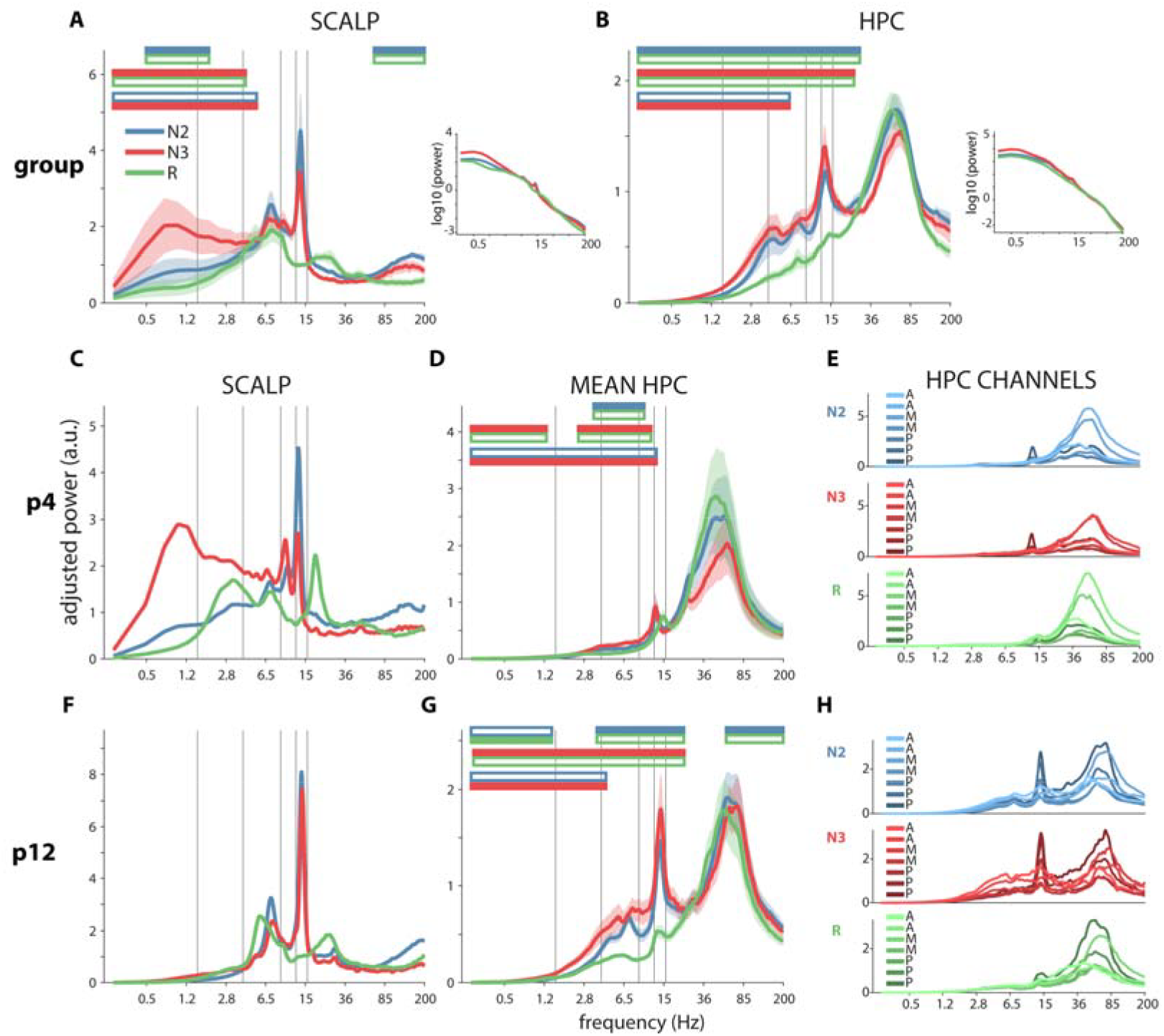
Scalp (Cz) and hippocampal sleep spectra. Top: Group-level spectra averaged across all 12 patients at the scalp (A) and in HPC (B). Error shading: standard error of the mean across patients. Horizontal color bars above plots indicate significant (P<0.05) pairwise stage differences (cluster-based permutation test). Filled color reflects stage with greater power. Gray vertical lines at 1.5, 4, 9, 12.5 and 16 Hz indicate approximate boundaries between SO, delta, theta, slow spindle, fast spindle, and faster activity. Right panels in (A) and (B) show same spectra without slope removal. The flattening of the raw spectrum at low (<4 Hz) frequencies results in comparatively low adjusted power after slope removal. Middle/bottom: Spectra of two example patients highlight differences between individuals, both at the scalp (C and F) and in HPC (D and G; error shading: standard error of the mean across data segments; stage differences as above). A further breakdown by HPC channel (E and H) reveals spectral differences along the longitudinal HPC axis (A: anterior [lighter color]; M: middle; P: posterior [darker color]; 7 HPC channels for both patients).

Because sleep spectra show substantial between-subject variability sleep (Cox et al., 2017; De Gennaro et al., 2005), we examined individual patients’ spectral profiles. While scalp NREM spectra commonly displayed strong peaks in the fast spindle range, the presence, prominence, and frequency of other spectral components varied (Fig. 2CF, Supp. Fig. 3). Theta and delta peaks, when present, could be largest in any sleep stage. Although slow spindles (9–12.5 Hz) are less prominent at central scalp positions (Zeitlhofer et al., 1997), several patients showed distinct peaks in the slow spindle range. This peak was clearly separate from nearby theta and fast spindle peaks in some instances (Fig. 2C), suggesting they reflect distinct oscillatory phenomena.

Considering HPC, channel-averaged spectra exhibited quite consistent gamma peaks across individuals, but otherwise showed large variability, particularly in the delta and theta range (Fig. 2DG, Supp. Fig. 3). A fast spindle peak could be discerned in NREM in most individuals, but was absent in some. Separate slow/fast spindle peaks were not observed. Also noteworthy is that while fast spindle peaks were absent during REM at the scalp (in line with sleep scoring criteria), such peaks were sometimes observed in HPC (Fig. 2DG), possibly related to asynchronous sleep stage transitions between brain structures (Durán et al., 2018; Emrick et al., 2016) and consistent with Fig. 1. In contrast, REM beta power frequently showed a peak at the scalp, but generally not in HPC.

Next, we considered spectra of single HPC channels for individual patients (Fig. 2EH). HPC channel spectra from a single patient were much more similar compared to spectra from different patients, with spectral peaks at similar frequencies. Nonetheless, peak amplitudes varied considerably along the longitudinal HPC axis. Power typically changed gradually from contact to contact, although this pattern was not always monotonic. Moreover, the direction of power increase (anterior-posterior or posterior-anterior) varied between individuals, and could occur in opposite directions for different frequency components in the same individual (e.g., N2/N3 spindle and gamma power in Fig. 2E), ruling out non-specific channel differences in signal amplitude. As a consequence, group-level comparisons between anterior, middle, and posterior HPC did not indicate systematic regional differences in spectral power (Supp. Fig. 4).

In sum, HPC spectral profiles varied with sleep stage, but this pattern differed substantially from that seen at the scalp. Moreover, we observed surprisingly diverse spectral patterns between individuals, and for HPC, between channels. Notably, this variability in the expression of spectral components is likely to affect their potential to engage in cross-frequency PAC, to which we turn next.

### 3.3 Single-channel PAC

We quantified two aspects of cross-frequency PAC. First, the strength of PAC (subsequently referred to as “coupling strength”, “coupling”, or simply “PAC”) signifies the degree to which activity of a faster frequency is non-uniformly distributed across the phase of a slower frequency. Second, coupling phase indicates the phase of the slower oscillation at which faster activity is preferentially expressed. Both metrics were assessed for every pair of frequencies in the 0.5–200 Hz range, separately for every one-minute artifact-free data segment, channel, sleep stage, and patient. Furthermore, coupling strengths for each segment were z-scored with respect to surrogate distributions. Note that our analytic approach does not distinguish between coupling stemming from two distinct oscillators, or from other phenomena (e.g., single asymmetric oscillator, harmonics; see Discussion). However, for the sake of readability we often describe effects in oscillatory terms (e.g., spindle-ripple coupling rather than spindle-band to ripple-band coupling). Given the large spectral variability reported in the previous section, we first sought to understand coupling dynamics for several example individual channels.

Assessing the presence of PAC for the same example subjects as in Fig. 2, both scalp (Fig. 3A) and HPC (Fig. 3D) channels showed prominent hotspots of frequency pairs where coupling strengths were significantly higher than zero (P<0.05, cluster-based permutation test across data segments). Note that a significant cluster may comprise more than one coupling phenomenon due to bridging between subclusters. Interestingly, while some cluster maxima coincided with peaks in the power spectrum, shown in the lower and right margin of each panel, other coupling effects were not accompanied by corresponding spectral peaks. For frequency pairs that were part of a significant cluster, we determined the mean coupling phase (Fig. 3BE) and, for several example frequency pairs, we evaluated the distribution of coupling phases across data segments (Fig. 3BE, insets). We now consider several observations in detail.

**Figure 3.**
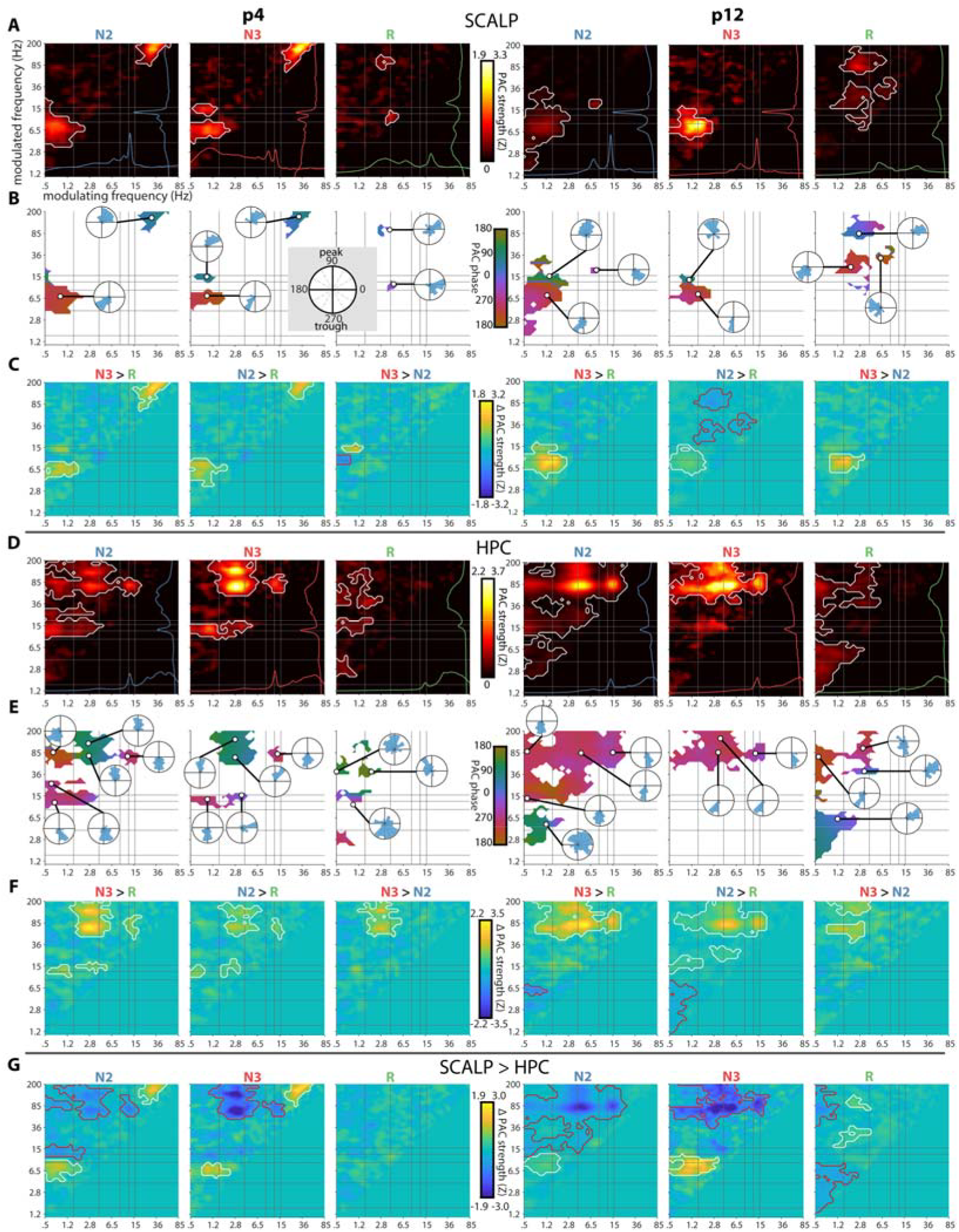
Phase-amplitude coupling for example channels. (A and D) Coupling strength at the scalp (A) and the most posterior HPC channel (D) for two example patients. White outlines indicate clusters of significantly higher than zero coupling across one-minute data segments (P<0.05, cluster-based permutation test). (B and E) Mean preferred coupling phases for significant frequency pairs. Insets: normalized circular histograms of coupling phase across data segments for example frequency pairs, indicating consistent phase preferences. Phases defined with respect to sine wave, with peak at 90° and trough at 270°. (C and F) Difference in coupling strength between sleep stages. White/red outlines indicate significant positive/negative clusters, respectively (P<0.05, cluster-based permutation test). (G) Difference in coupling strength between scalp and HPC. White/red outlines indicate significant positive/negative clusters, respectively (P<0.05, cluster-based permutation test).

Scalp PAC in both example patients was most clearly seen during NREM between SO and theta and between SO and fast spindle activity (Fig. 3A), with phases of maximal coupling towards the SO trough and peak, respectively (Fig. 3B), consistent with previous findings (Cox et al., 2018; Gonzalez et al., 2018; Mölle et al., 2011). Note that theta clusters extended into the slow spindle range without clear boundaries (see section 4.4). Coupling in these ranges was significantly stronger in NREM than REM, and also differed between N3 and N2 (Fig. 3C; P<0.05, cluster-based permutation tests), indicating the stage-specificity of these effects. Scalp PAC during REM was generally less pronounced, although there was evidence for PAC between delta/theta and beta/gamma in both patients. Other effects differed between patients. For example, patient p12 exhibited N2 theta–beta coupling coinciding with a theta spectral peak (Fig. 3A, right). In contrast, patient p4 demonstrated strong PAC during NREM between beta/gamma and >150 Hz activity (Fig. 3A, left), likely reflecting muscle activity (see section 4.1).

HPC PAC was also clearest during NREM. Both example patients expressed strong evidence for the modulation of high gamma activity, with distinct clusters centered at ~80 Hz and ~140 Hz, consistent with ripple activity (Fig. 3DE). Ripple activity was coupled to the oscillatory phase of various slower frequency bands, including the SO, delta, theta, and fast spindle ranges, depending on NREM stage and patient. Note that ripple locking to the SO trough, rather than the peak, suggests an opposite polarity compared to the scalp, with the SO trough reflecting the physiological UP state for these channels (Nir et al., 2011). The observation of delta–ripple coupling is consistent both with sharp wave-ripple sequences (Oliva et al., 2018), and with ripple coupling to a delta rhythm of different origin. Interestingly, whereas ripple activity was locked to the trough of both the SO and delta ranges for patient p12 (Fig. 3D, right), it was tied to opposite phases for SO (trough) and delta (peak) activity for patient p4 (Fig. 3D, left). Distinct phase preferences for SO and delta activity fit the idea that these slow components reflect separate phenomena (Amzica and Steriade, 1998). Spindle–ripple coupling was tied to the spindle trough, as reported previously (Clemens et al., 2011; Staresina et al., 2015). Direct stage comparisons indicated that ripple activity was most strongly coupled to slower activity during N3, followed by N2, and finally REM (Fig. 3F). In addition to the modulation of ripple activity, spindle activity was preferentially organized into the SO/delta trough in both patients (Fig. 3DE), again consistent with opposite polarities in HPC and scalp, although coupling was weak for p12. N2 SO– delta/theta coupling appeared in patient p12 (Fig. 3DE, right), but with a phase preference opposite to that seen at the scalp. The pattern of HPC PAC during REM was less clear, although some coordination of faster activity by the SO/delta phase was present, which could even surpass that seen during NREM (e.g., N2/N3 SO– theta in Fig. 3D, right). Finally, direct contrasts between HPC and scalp underscore the regional specificity of many of the aforementioned effects, including stronger N2/N3 ripple and N2 spindle modulation in HPC, but enhanced N2/N3 theta modulation at the scalp (Fig. 3G).

Combined, these results illustrate clear temporal coupling between various pairs of oscillatory rhythms. Although the expected coordination among SO, spindle, and ripple rhythms was present, coupling extended well beyond these frequency ranges, with different dynamics for HPC and scalp, and different patterns for NREM and REM sleep. Moreover, coupling dynamics differed between patients, potentially related to corresponding individual differences in spectral profiles.

### 3.4 Single-patient PAC along the hippocampal axis

Next, we considered PAC for all channels along the HPC axis within individuals. Fig. 4AB shows the complete profile for two example patients (additional example in Supp. Fig. 5), including patient p4 described in the previous section. Although coupling patterns showed a certain degree of similarity across channels for both patients, clear differences were also present, such that frequency pairs showing large and above-chance coupling on one channel could show much reduced (and non-significant) coupling on other channels. Interestingly, while the degree of SO– spindle and spindle–ripple coupling corresponded to the height of the spindle peak in the spectrum for patient p4, this phenomenon was not observed for patient p11. (Also note the absence of clearly distinguishable spindle–ripple clusters in the latter patient.) In contrast, channels with largest ripple power were generally not the ones that were modulated most strongly by slower rhythms.

**Figure 4.**
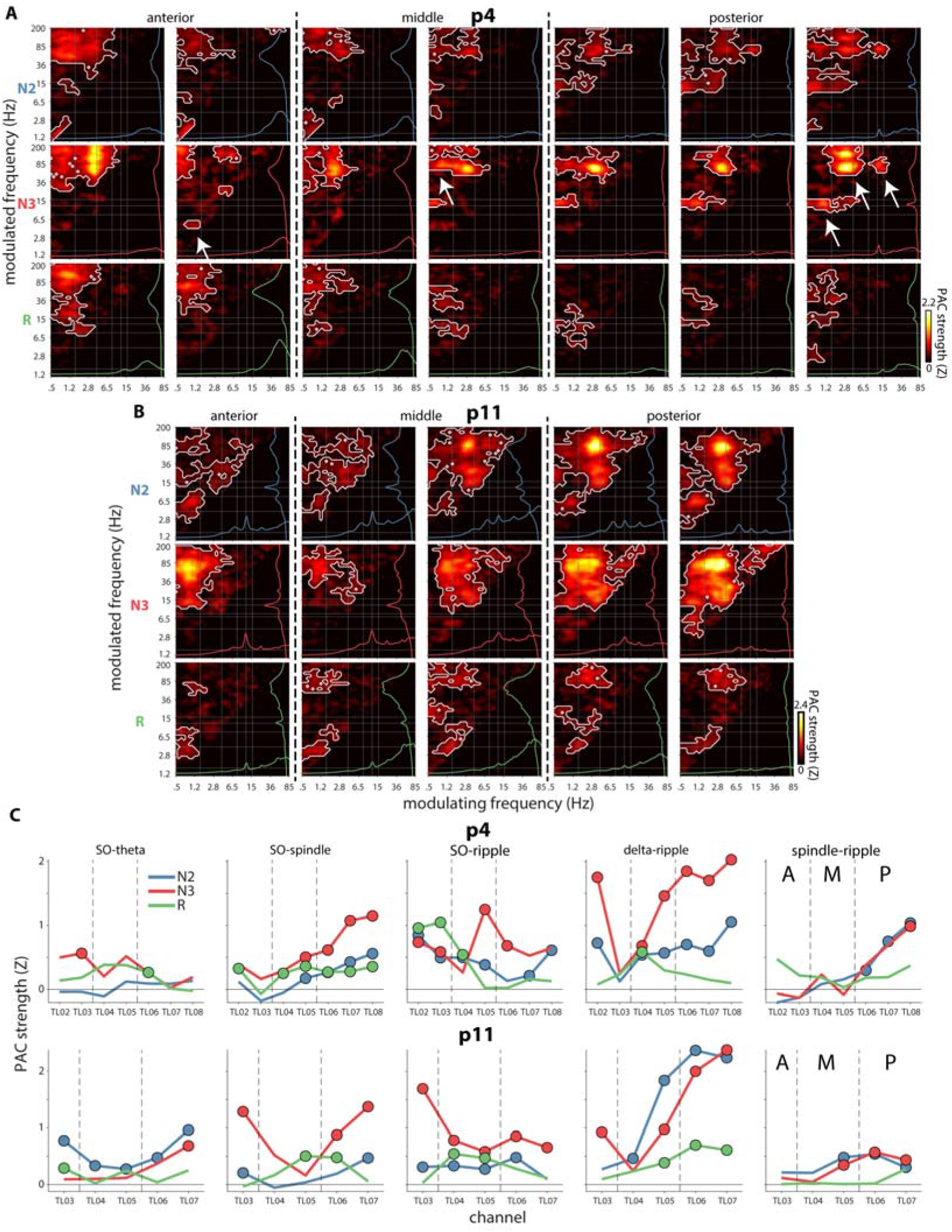
Phase-amplitude coupling along hippocampal axis for example patients. (A and B) Coupling strengths for all HPC contacts in two example patients. White outlines indicate clusters of significantly higher than zero coupling (P<0.05, cluster-based permutation test). (C) Coupling strengths for five frequency pairs, indicated in (A) with arrows. Presence of marker indicates frequency pair is part of significant cluster in (A) or (B). Dashed lines separate anterior (A), middle (M), and posterior (P) channels.

Interestingly, patients expressed distinct longitudinal profiles of HPC coupling, such that frequency pairs could be most strongly coupled at different HPC sites for different patients. These findings echo our earlier observations for spectral power (Fig. 2). To further examine longitudinal coupling profiles, we plotted coupling strengths along the HPC axis for five example frequency pairs (Fig. 4C). These plots not only illustrate distinct regional coupling patterns for the two patients, but also distinct patterns for different frequency pairs and sleep stages. Note the different profiles for SO–ripple and delta–ripple coupling, again suggesting that SOs and delta activity constitute distinct phenomena. Also noteworthy is that while coupling was generally weakest during REM, REM coupling was nonetheless apparent for most frequency pairs, and surpassed the level seen during NREM on some channels (e.g., anterior SO–ripple coupling for patient p4). The latter finding fits with the presence of SOs during REM sleep (Funk et al., 2016).

We note that although longitudinal profiles of Fig. 4 show a modest degree of resemblance between patients (e.g., stronger delta-ripple and spindle-ripple PAC in posterior HPC), these observations do not imply systematic effects. Indeed, another example patient showed strong delta-ripple coupling along the entire HPC axis, whereas reliable NREM spindle-ripple was not observed at all (Supp. Fig. 5). Group-level evaluation of PAC in hippocampal subregions will be presented in section 3.6.

In sum, while oscillatory interactions can be seen along the entire axis of human HPC, whether and how strongly specific frequency pairs interact varies from contact to contact, consistent with the notion of spatially localized sleep oscillations (Funk et al., 2016; Nir et al., 2011; Vyazovskiy et al., 2011). Moreover, HPC coupling profiles vary considerably between individuals, extending similar observations from non-invasive approaches (Cox et al., 2018, 2017; De Gennaro et al., 2005).

### 3.5 Group-level PAC

We next examined scalp and HPC PAC across patients. As shown in Fig. 5A (left), scalp coupling was most evident during N3 between the SO and the theta/spindle bands, with a similar, albeit weaker, cluster present during N2. In contrast, REM showed evidence of weak coupling between delta/theta phase and beta/gamma power, consistent with the effects of individual patients (Fig. 3A). Both NREM stages also exhibited a cluster of gamma–high gamma coupling (maximum at 40 × 200 Hz), similar to that seen for one of the example patients (Fig. 3A). This finding is likely due to muscle rather than neural activity (see section 4.1). Direct stage comparisons indicated significantly stronger PAC in N3 than REM for the SO-theta effect (Fig. 5B, left). Overall, these findings corroborate previous scalp findings (Cox et al., 2018; R. Cox et al., 2014; Mölle et al., 2011), while also indicating the consistency of scalp-measured PAC across individuals.

**Figure 5.**
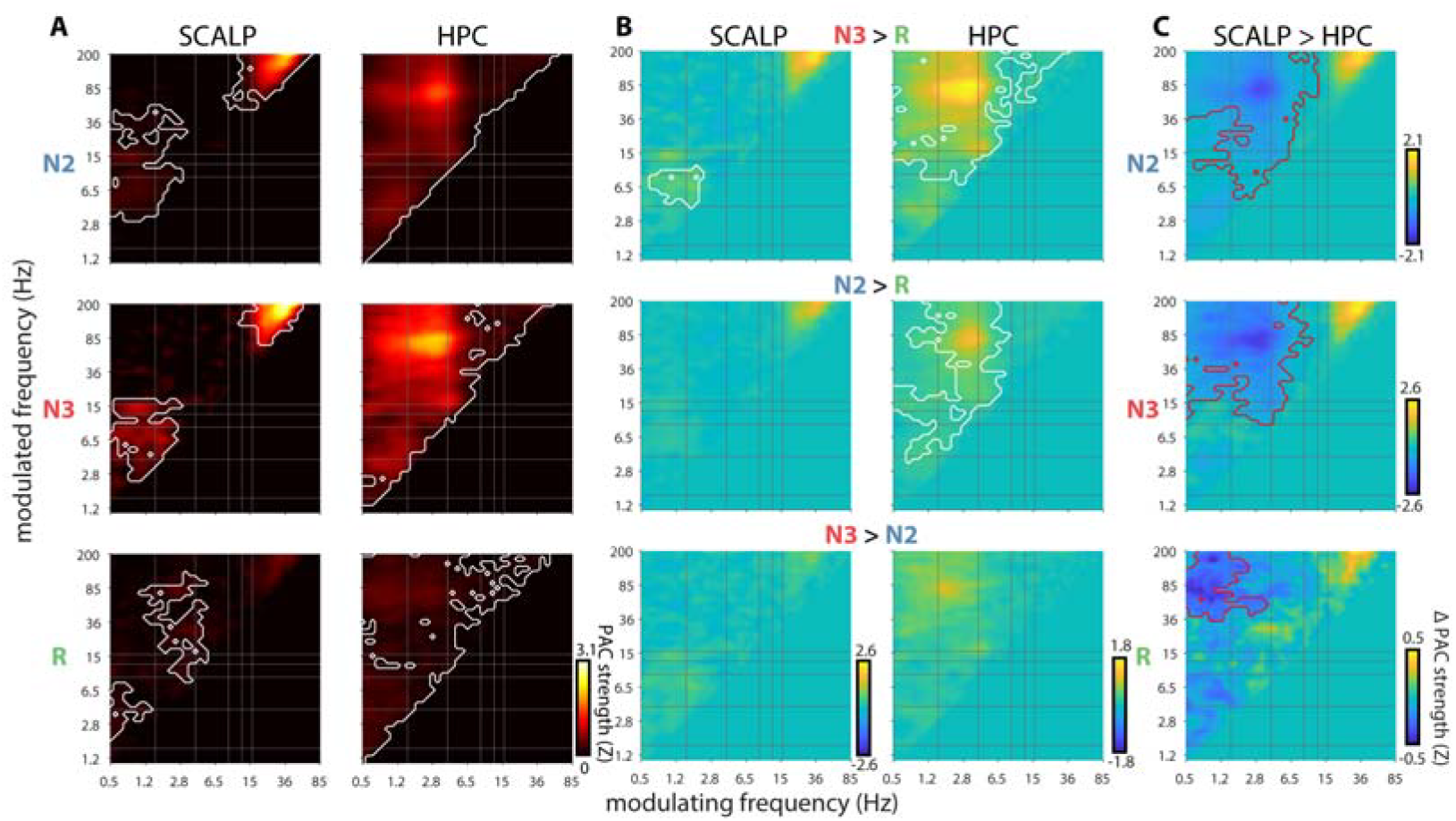
Group-level phase-amplitude coupling. (A) Mean coupling strength for scalp (left) and HPC (right). White outlines indicate clusters of significantly greater than zero coupling (P<0.05, cluster-based permutation test). (B) Mean difference in coupling strength between sleep stages for scalp (left) and HPC (right). (C) Difference in coupling strength between scalp and HPC. White/red outlines in (B) and (C) indicate significant positive/negative clusters, respectively (P<0.05, cluster-based permutation test).

For HPC, coupling profiles were first averaged across available HPC contacts for each patient. Subsequent group analyses revealed prominent hotspots of delta– ripple coupling in both N2 and N3 (Fig. 5A, right). In contrast, clearly delineated foci for other coupling phenomena that might be expected (e.g., SO–spindle, SO–theta, spindle–ripple) were not apparent at the group level. REM PAC was much lower, showing weak hotspots in the SO–ripple and delta–ripple ranges. Nonetheless, statistical evaluation signaled massive clusters for every sleep stage, essentially comprising all frequency pairs. (An alternative statistical approach using False Discovery Rate-based (Benjamini and Hochberg, 1995) rather than cluster-based correction yielded similar results (not shown)). Note that this observation should not be taken to imply that every frequency is consistently modulated by every other frequency. Rather, these findings suggest that the aforementioned variability in coupling dynamics between patients and individual HPC channels results in above-zero group averages for every frequency pair, which then form a single big cluster. Direct stage comparisons indicated stronger coupling in both NREM stages relative to REM, for frequency pairs including, but not limited to, the delta–ripple range (Fig. 5B, right). Directly contrasting scalp and HPC coupling profiles revealed significantly stronger PAC in HPC for each stage, mainly between SO/delta and beta/gamma frequencies (Fig. 5C).

While the group analyses of Fig. 5A provide evidence for the general presence of PAC, they offer limited information about which exact frequency pairs are coupled, particularly in HPC. To obtain a qualitative view how often particular frequency pairs were coupled in our sample, we adopted an unsupervised clustering approach. Specifically, for every subject, sleep stage, and channel, we extracted all frequency pairs showing a local maximum in coupling strength. Of these, the frequency pairs corresponding to coupling strengths above a threshold were submitted to a density-based clustering algorithm (Ester et al., 1996). We employed both adaptive, data-derived thresholds (i.e., different thresholds for every subject, sleep stage, and region) and fixed thresholds (PAC > 2). This yielded, for every subject, sleep stage, and brain region, a relatively small number of clusters, whose centers were taken as frequency pairs expressing coupling. Clustering results for several example patients using adaptive thresholds are shown in Supp. Fig. 6.

Fig. 6 aggregates the clustering algorithm results across individuals as heat maps, with color indicating how many subjects exhibited evidence of coupling at each frequency pair, using either adaptive (Fig. 6A) or fixed (Fig. 6B) thresholds. Additional plots below and to the right of each panel indicate how many unique individuals showed coupling for every modulating and modulated frequency, respectively.

**Figure 6.**
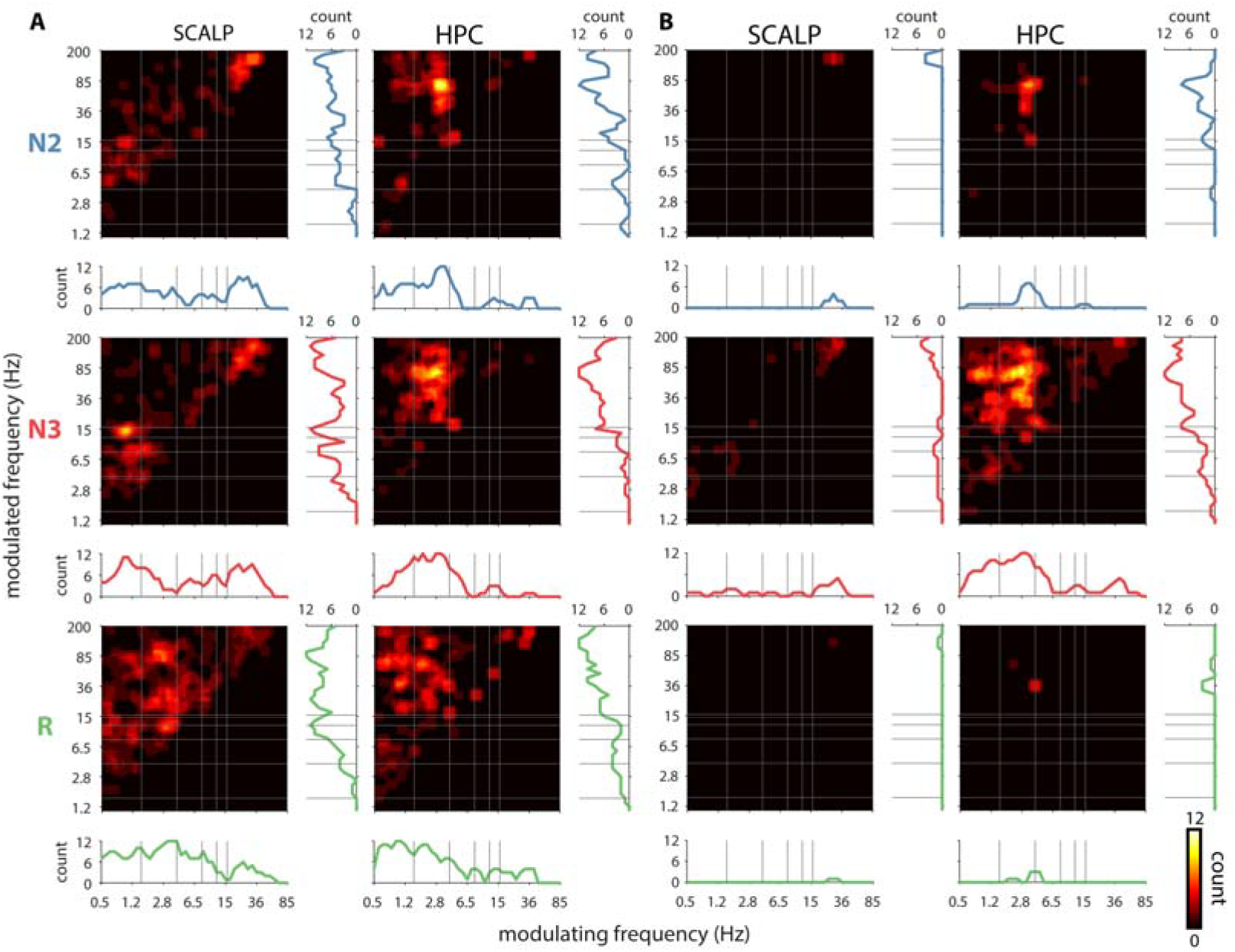
Group-level counts of phase-amplitude coupling. Number of individuals showing PAC for each frequency pair using adaptive (A) and fixed (B) thresholds at the scalp (left) and HPC (right) for different sleep stages (rows). Line plots indicate number of unique individuals showing PAC at each modulating (bottom) or modulated frequency (right) bin.

Using adaptive thresholds, scalp NREM showed strongest group effects for SO–spindle, SO–theta, and gamma–high gamma frequency pairs (Fig. 6A, left), similar to the effects using conventional group statistics in Fig. 5A (left). Scalp REM showed most consistent evidence for coupling between delta/theta modulating frequencies and spindle/beta/ripple modulated frequencies. In contrast, the clustering approach yielded a more clearly defined NREM profile for HPC (Fig. 6A, right), compared to the diffuse appearance of the conventional group approach (Fig. 5A, right), whereas HPC REM coupling continued to show a highly variable pattern (detailed description of coupling effects in next paragraph). However, it is important to note that overall coupling strengths were much lower in REM compared to NREM, and reduced at the scalp compared to HPC. Consequently, the fixed-threshold approach abolished the majority of scalp effects, and for REM, only a cluster of delta/theta–beta coupling remained in HPC (Fig. 6B).

Considering the adaptive threshold approach in detail, NREM scalp coupling was most consistently seen for N3 SO–spindle (9/12 individuals) frequency pairs, followed by N2 SO–spindle (5), N3 SO–theta (5), N2 SO–theta (4), N2 SO–theta/slow spindle (4), and N3 SO/delta (4) frequency pairs. In HPC, NREM coupling was highly consistent for N2 (12) and N3 (11) delta–ripple frequencies, followed by coupling between modulating frequencies on the delta/theta border and their modulated frequencies covering the high spindle/low beta range (N3: 5; N2: 4). SO–ripple coupling was seen in both N2 and N3 (4), whereas SO–delta coupling only occurred in N2 (4). Spindle–ripple coupling emerged in only three patients in both NREM stages, whereas above-threshold SO–spindle PAC was not observed at all. A complete overview of all NREM effects present in at least 3 subjects (25%) can be found in Supp. Table 2. The even larger variability of REM coupling profiles, due to the inclusion of weaker (and likely noisier) effects, precludes meaningful construction of a similar table for REM.

In sum, while sleep oscillations are relatively consistently coupled at the scalp, group-level PAC within HPC was much less systematic, with many effects occurring in only a minority of individuals, further confirming the observations from individual patients (Fig. 3 and 4).

### 3.6 Group-level PAC in hippocampal regions

Finally, we considered group-level PAC for anterior, middle, and posterior HPC regions. Interestingly, direct comparisons between HPC regions revealed two systematic differences (Fig. 7). First, N2 coupling between SOs and activity in the theta/spindle range was significantly stronger in anterior than middle HPC. Second, SO/delta–ripple PAC showed a difference during REM, with greater coupling in anterior HPC relative to both middle and posterior HPC. (This effect was also present at the subject-level for patient p4 in Fig. 4C.) Note that both effects occurred in the absence of systematic regional differences in spectral power (Supp. Fig. 4). Region-specific analyses considering the presence of coupling (Supp. Fig. 7) and sleep stage differences (Supp. Fig. 8) yielded results similar to those seen across all HPC channels, as presented in Fig. 5A (right) and Fig. 5B (right), respectively.

## 4 Discussion

The present study offers a comprehensive analysis of cross-frequency phase-amplitude coupling at the scalp and in the hippocampus of sleeping humans. Considering frequency pairs over an extensive 0.5–200 Hz range, we found strong evidence for the presence of spectral interactions, with the overall strength of coupling greater in HPC than at the scalp, and greater during NREM than REM sleep. Scalp PAC during NREM consisted of SO–theta and SO–spindle coupling, while theta– gamma coupling was seen during REM. In contrast, HPC PAC was seen for many frequency pairs, including both expected (e.g., NREM SO–ripple, SO–spindle, delta– ripple, spindle–ripple), and novel interactions (e.g., NREM SO–delta, SO–theta, REM delta/theta–beta). At the same time, coupling profiles differed substantially between individuals, and between HPC sites within individuals.

### 4.1 Sleep oscillations at the scalp

Scalp-level group analyses indicated well-known NREM coupling phenomena between SO phase and amplitudes of both spindle and theta activity (Fig. 5A) (Cox et al., 2018; R. Cox et al., 2014; Gonzalez et al., 2018; Klinzing et al., 2016; Mölle et al., 2011). These effects were stronger in N3 compared to N2, consistent with the lower density of SOs in N2 (Cox et al., 2018) (likely including a large proportion of isolated K-complexes). Although individuals expressed variability in both power spectra (Fig. 2CF, Supp. Fig. 3) (Cox et al., 2017; De Gennaro et al., 2005) and coupling profiles (Fig. 3A), SO–theta and SO–spindle effects were relatively consistent across individuals, thus leading to circumscribed effects at the group level. Importantly, this corroboration of known effects establishes the general suitability of our analytical approach.

Interestingly, while coupling was generally weaker during REM, we observed significant coupling in this sleep stage between the phase of the delta/theta rhythm and amplitudes in the beta/gamma range, consistent with the presence of theta and beta peaks in the power spectrum. We are not aware of previous studies reporting such REM coupling in human scalp EEG, although this finding fits with similar findings in rodents (Brankačk et al., 2012; Scheffzük et al., 2011) and human parahippocampal cortex (Clemens et al., 2009) (also see section 4.2).

During NREM, clusters of beta/gamma–high gamma (>150 Hz) coupling were seen at both the group and individual levels. However, high-frequency (>~80 Hz) components of non-invasive EEG cannot be assumed to reflect neural activity without additional validation. In contrast, scalp-recorded electromyographic activity extends well beyond this range (Muthukumaraswamy, 2013). For this reason, and because the observed coupling effect was absent during the atonic state of REM sleep (Fig. 5A), along with a REM-related reduction in high-gamma power (Fig. 2A), the most likely explanation for this effect is muscle activity. Indeed, coupling of high-frequency muscle activity to lower frequency scalp oscillations has previously been reported (So et al., 2016; Yang et al., 2016), but it is presently unclear whether our finding of beta/gamma–high gamma PAC represents such a case.

Recent rodent work indicates that spindle oscillations coordinate the expression of ripple-like (~100 and ~200 Hz) activity in neocortex (Averkin et al., 2016). However, our scalp recordings did not show modulation of frequencies >85 Hz by NREM spindles or SOs, in line with previous scalp analyses (Staresina et al., 2015). Similarly, scalp spectra were not indicative of obvious ripple activity (Fig. 2ACF), suggesting that the possibility that ripples can be non-invasively recorded from the scalp is limited (Mooij et al., 2018, 2017).

A 9–12.5 Hz slow spindle peak was observed in only a minority of cases, likely due to suboptimal positioning of the central scalp electrode relative to the predominantly frontal generators of slow spindles (Cox et al., 2017; Zeitlhofer et al., 1997). As a consequence of this, as well as the limited spectral precision of our methods (see section 4.4), our PAC findings do not directly speak to the recently raised possibility that theta waves are mistaken for slow spindles (Gonzalez et al., 2018). However, we note that distinct slow spindle and theta components were sometimes apparent in the power spectrum (Fig. 2C).

### 4.2 Sleep oscillations in the hippocampus

Considering HPC, we found strong evidence for various oscillatory interactions, particularly during NREM. Yet, coupling patterns were strikingly diverse across individuals and HPC contacts. Furthermore, the overall profile of oscillatory pairs engaging in HPC PAC deviated substantially from the one seen at the scalp, indicating the regional specificity of oscillatory organization.

Across individuals and channels, NREM delta-ripple PAC was the clearest and most consistent coupling phenomenon (Fig. 4, 4A, 5), corroborating earlier reports (Axmacher et al., 2008; Staresina et al., 2015). A likely possibility is that the delta aspect of this coupling effect reflects the sharp wave component of sharp wave– ripple complexes, since the spectral representation of sharp waves peaks in the delta range (but extending up to 10 Hz) (Oliva et al., 2018). Also noteworthy in this respect is the observation of delta–beta (20–25 Hz) coupling in several instances. This beta component could be akin to the “slow gamma” band modulation recently shown to constitute the rhythmicity of overlapping ripples (Oliva et al., 2018). That said, the current findings do not rule out the possibility of alternative, non-sharp wave, delta oscillations contributing to delta-ripple PAC.

In this light, it is important to point out that HPC delta–ripple and SO–ripple coupling likely constitute distinct phenomena. First, above-threshold SO–ripple coupling appeared in only a third of our sample, compared to the full sample for delta–ripple PAC. Note that this observation corresponds to a previous report (partially overlapping with the data presented in the current paper) of ripples being most strongly modulated by delta rather than SO activity (Staresina et al., 2015). Second, longitudinal profiles for these frequency pairs differed substantially within individuals (Fig. 4C). Third, separate SO–ripple and delta–ripple hotspots could be discerned within an individual, sometimes with opposite phase preferences (e.g., Fig. 3D, left, N2). Fourth, we observed SO and delta activity to be coupled themselves in several instances, similar to earlier findings in neocortex (Steriade et al., 1993). Overall, these findings fit earlier arguments that <4 Hz activity does not constitute a single phenomenon, but may be separated into physiologically and functionally distinct entities (Achermann and Borbély, 1997; Amzica and Steriade, 1998; Benoit et al., 2000).

Also noteworthy is the observation of N2 coupling in HPC between SOs and the high theta/slow spindle range (Fig. 4B, 5A). Given that we did not observe separate slow and fast spindle peaks in HPC, we tentatively interpret the modulated frequency to reflect theta activity. Recent non-invasive findings indicate a role for SO–theta PAC in memory consolidation (Schreiner et al., 2018). The presence of a similar dynamic in HPC lends further support to this idea, although it is striking that this phenomenon was not observed in N3, during which it was most prominently seen at the scalp.

While we observed unambiguous clusters of strong SO–spindle and spindle– ripple coupling for individual HPC channels (Fig. 3 and 4), these findings did not emerge consistently at the group level, using either standard (Fig. 5) or unsupervised clustering approaches (Fig. 6). This finding is somewhat surprising in light of recent demonstrations of SO–spindle–ripple coupling in human (para)hippocampus (Clemens et al., 2011, 2007; Staresina et al., 2015). However, an important methodological difference is that these previous studies mostly limited PAC calculations to detected waveforms, whereas we considered continuous data without imposing specific detection criteria (see section 4.4). Moreover, the study reporting hierarchical nesting of all three waveforms (partially overlapping with the current data) (Staresina et al., 2015) included only the contact with highest spindle power, thereby making analyses contingent on the presence of clear spindles. Indeed, while we observed spindle spectral peaks on at least one HPC channel in most individuals, many channels were not suggestive of spindle activity (see spectral profiles inside panels of Fig. 4AB). This may also explain inconsistent reports regarding the presence of spectral spindle peaks in (para)hippocampus (Montplaisir et al., 1981; Nakabayashi et al., 2001). More generally, although absolute signal amplitudes are higher in HPC, both SOs and spindles are much less visually apparent in HPC compared to the scalp. In sum, while our results confirm the existence of SO– spindle and spindle–ripple coupling in human HPC, our findings also indicate that this coupling may not be as ubiquitous as previously thought.

Similar to scalp findings, hippocampal coupling was much reduced during REM compared to NREM (Fig. 5AB). While conventional group analyses did not indicate clearly defined REM coupling, unsupervised clustering suggested the presence of delta/theta–beta (4 vs. 36 Hz) coupling in three patients (fixed thresholds, Fig. 6B). This result is reminiscent of previous findings in human parahippocampal cortex (Clemens et al., 2009), although in that report the modulating frequency was slower (1–3 Hz) and modulation was strongest in the 60– 100 Hz range. As such, our findings do little to answer whether the human functional analog of rodent hippocampal theta lies in the SO/delta (Bódizs et al., 2001; Clemens et al., 2009) or theta (Cantero et al., 2003) range, since our 4 Hz modulating frequency falls right in between. Rather, our observations of variable spectral and coupling profiles during REM suggest that both SO/delta and theta frequencies can be observed (also see section 4.4). More generally, the ambiguous evidence for REM theta-like activity in human HPC underscores the need to consider species differences in oscillatory organization. Indeed, HPC theta activity was typically more pronounced in NREM than REM (Fig. 2). Also noteworthy is that SO–ripple coupling was found to be stronger in HPC vs. scalp in all sleep stages including REM (Fig. 5C), suggesting that SOs may also occur in this sleep stage (Funk et al., 2016) (also see section 4.3). These findings are consistent with asynchronous sleep stage transitions between scalp and HPC (Durán et al., 2018; Emrick et al., 2016), as further suggested by REM spindle occurrences within HPC (Fig. 1).

### 4.3 Regional and individual variability in hippocampus

A key observation of the present study is the large heterogeneity in oscillatory and coupling profiles between HPC contacts and individuals. What could underlie this variability? First, several technical factors could play a role. For example, electrode orientation relative to generating fields could render contacts differentially sensitive to certain waveforms. Related, HPC subfields (e.g., CA3, CA1, DG) are known to participate differently in sleep-related network activity (Isomura et al., 2006; Oliva et al., 2018). Unfortunately, anatomical resolution in the current study cannot answer whether subfield identity could account for our findings. It also deserves mention that the observed regional variability makes it unlikely that HPC findings are driven by the mastoids reference, since such an influence should impact all HPC contacts equally.

While the preceding factors essentially argue that heterogeneity is a rather uninteresting consequence of data acquisition, another possibility is that the observed differences constitute true variability. Specifically, regional HPC variability is consistent with accumulating evidence of the local nature of sleep oscillations (Cox et al., 2018; R. Cox et al., 2014; Nir et al., 2011; Vyazovskiy et al., 2011). Particularly telling in this respect is that different spectral components could be strongest at opposite ends of HPC for the same individual (e.g., N2 spindle and ripple activity in Fig. 2E). Similarly, distinct coupling phenomena could be most pronounced at different contacts (e.g., N3 SO–ripple in anterior, and delta–ripple in posterior HPC; Fig. 4C).

At the group-level, HPC regional variability emerged as significantly enhanced SO–theta and SO–spindle PAC in anterior vs. middle HPC during N2, as well as enhanced SO–ripple coupling in anterior vs. middle/posterior HPC during REM (Fig. 7). However, these effects should be interpreted with caution since these frequency pairs did not emerge as clearly defined clusters when assessing the presence of coupling (Fig. 5 and 5, Supp. Fig. 7). Nonetheless, these regional differences provide some evidence that the pertaining frequency bands were in fact coupled. Furthermore, regional variability in SO–ripple coupling during REM suggests that aspects of NREM electrophysiology can also be observed during REM in HPC (also see section 4.2).

**Figure 7.**
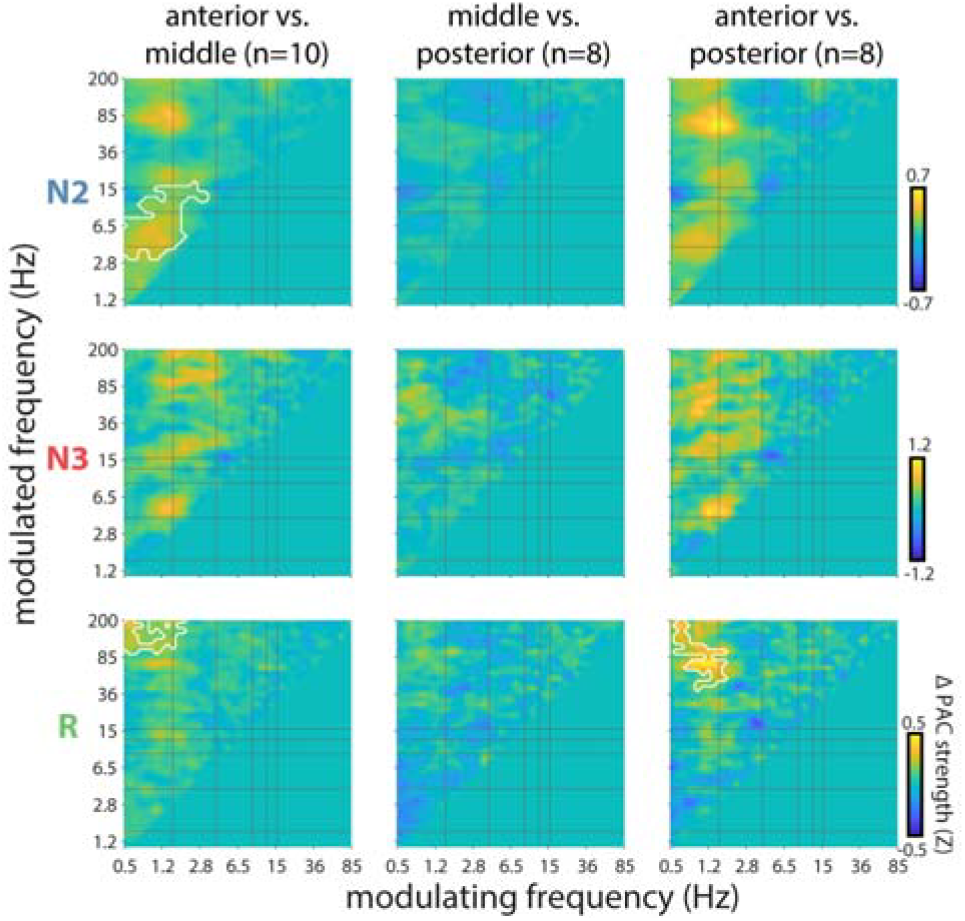
Regional differences in hippocampal phase-amplitude coupling. Mean difference in coupling strength between anterior and middle, middle and posterior, and anterior and posterior HPC for each sleep stage. White outlines indicate significant positive clusters (P<0.05, cluster-based permutation test; no negative clusters present). Note different sample sizes for each regional comparison due to variable electrode coverage.

In contrast, no systematic regional coupling differences were observed for SO–ripple, delta–ripple, or spindle–ripple pairs during NREM, nor for SO–spindle coupling during N3. Similarly, we did not observe any systematic regional differences in spectral power for any frequency band during any sleep stage (Supp. Fig. 4). However, we note that the anatomy-based division into HPC thirds does not necessarily result in functionally homologous subregions across patients. Thus, defining regions functionally might have yielded different results. Although we did not have specific predictions regarding oscillatory structure in anterior, middle, and posterior HPC, we examined this possibility because of well-known intrinsic and extrinsic wiring differences between anterior and posterior HPC (or the analogous ventral/dorsal divide in rodent HPC) (Fanselow and Dong, 2010; Strange et al., 2014), and evidence of differential regional relations with human memory (Bonnici et al., 2012; Dandolo and Schwabe, 2018; Ludowig et al., 2008; Poppenk and Moscovitch, 2011). In sum, with the potential exception of N2 SO–spindle coupling, we conclude that the sleep oscillations commonly thought relevant for memory consolidation (i.e., SOs, spindles, ripples), are not differentially present or coupled along the hippocampal axis in a systematic fashion.

Nonetheless, we emphasize that this group-level conclusion does not reflect individual patients, who showed marked differences in spectral power and PAC along the HPC axis. We suggest that individual differences in HPC PAC constitute true variability representative of the population, similar to replicable individual differences of scalp sleep oscillations (Cox et al., 2018, 2017; De Gennaro et al., 2005; Massimini et al., 2004; Rusterholz et al., 2018). Whether such variability is further related to general (e.g., cognitive functioning, age) or disease-related (e.g., epilepsy history, medication) factors cannot be determined from the present sample.

### 4.4 Methodological considerations

Although findings from medicated epilepsy patients necessarily warrant caution when generalizing to healthy populations, sleep architecture (Table 1) and scalp power spectra (Fig. 2A, Supp. Fig. 3) were within normal ranges, only data from the non-pathological HPC was used, and a stringent artifact rejection procedure removed >50% of data, making it unlikely that our results are due to epileptiform activity. However, we point out the possibility that delta-ripple PAC partly reflects pathological spike-ripple coupling (Weiss et al., 2016).

While our sample size (N=12) was sufficiently large to allow group-level inferences, its modest size, though not atypical for invasive studies, has implications for generalizability. In particular, while the general notion of individual and regional variability may be extrapolated to the population as a whole, the specific pattern of variability should not. For example, while NREM SO–theta, SO–spindle, delta–ripple, and spindle–ripple coupling were all observed, the present sample did not contain a single individual/channel clearly expressing all of these effects. Nonetheless, it is not unreasonable to expect the co-existence of each of these effects in the larger population. Similarly, the population likely harbors individuals with idiosyncratic spectral peaks and coupling effects for frequencies not seen in the present sample. In this light, we also note that previous invasive studies may have reached quite different conclusions (e.g., presence vs. absence of spectral spindle peaks; SO/delta vs. theta activity during REM) due to taking small samples from a highly variable population, thus increasing the likelihood of capitalizing on chance.

In the present approach, PAC was calculated over continuous data. This contrasts with discrete approaches where PAC calculations are contingent on the presence of specific waveforms, detected according to specific criteria of frequency, amplitude, and/or duration. In line with our objective of assessing spectral interactions in a data-driven fashion without setting *a priori* restrictions on the forms of coupling that might be present, we favored the continuous approach. We also note that the employed wavelet approach is inherently limited in spectral (and temporal) resolution, hampering isolation of closely spaced spectral components (e.g., theta, slow spindle, and fast spindle components). Moreover, the use of a fixed set of wavelet frequencies does not allow subject-specific targeting of oscillatory phenomena that vary in frequency between individuals (Cox et al., 2017; Ujma et al., 2015). However, given that our approach identified various previously described coupling phenomena, we believe these methodological choices do not pose a major concern.

Metrics of PAC are typically sensitive to both genuine cross-frequency coupling and asymmetrical features of a single oscillator (Cole and Voytek, 2017). However, specific and distinct generating circuits can reasonably be ascribed to several spectral components we found to be coupled (e.g., SO, spindle, ripple, sharp wave). Also worth mentioning is that the present PAC metric is not sensitive to multimodal phase preferences, for which other approaches may be preferred (Tort et al., 2010). While inspections of raw traces indicated the presence of oscillatory activity in various frequency bands (Fig. 1, Supp. Fig. 1), it remains a possibility that some of our reported coupling effects are at least partly due to factors other than two separate oscillators. Ultimately, detailed waveform analyses are required to fully understand the origin of each of the observed effects (Cole and Voytek, 2017), particularly those that have little precedence in the literature.

In several instances we observed that the presence of coupling coincided with corresponding spectral peaks, and that coupling strength scaled with the height of the modulating peak (e.g., N3 spindle–ripple PAC in Fig. 4A). In other instances, this relation was not apparent (e.g., N3 spindle–ripple PAC in Fig. 4B), indicating no straightforward mapping between power and PAC. Moreover, it has been suggested that the presence of a narrowband spectral peak for the modulating frequency is required to meaningfully interpret PAC (Aru et al., 2015). However, the most robust coupling phenomenon in HPC, delta–ripple PAC, occurred without clear peaks in the delta range (Fig. 2B). Finally, there were instances where two frequencies showed clear spectral peaks, but were not significantly coupled, indicating that coupling does not invariably emerge when multiple oscillators are present.

### 4.5 Functional implications

The present findings are relevant to theories on the role of coupled sleep oscillations in memory consolidation. HPC PAC was seen for a remarkably diverse range of frequency pairs during NREM, with coupling phenomena extending well beyond previously described interactions between SOs, spindles, and ripples (Clemens et al., 2011, 2007; Staresina et al., 2015). Indeed, these classical coupling patterns were less consistently present than we expected, whereas delta–ripple PAC was reliably present in the full sample. That said, SOs, spindles, and ripples are robustly tied to plasticity and consolidation processes (Axmacher et al., 2008; Cairney et al., 2018; R. Cox et al., 2014; Huber et al., 2004; Latchoumane et al., 2017; Niethard et al., 2018; Schönauer et al., 2017; Zhang et al., 2018), whereas the functional relevance for other coupling pairs remains to be demonstrated. It is also possible that the coordination of HPC ripple and spindle oscillations by spindles and SOs, respectively, becomes more evident when considering the phase of neocortical rather than HPC spindle/SO activity. Such cross-regional spectral interactions have been described in animals (Siapas and Wilson, 1998; Sirota et al., 2003) and humans (Staresina et al., 2015), but were outside the scope of the present study. In contrast to NREM, the functional relevance of REM sleep remains enigmatic (Rasch and Born, 2013; Stickgold and Walker, 2013). Whatever its function, our findings indicate that the role of coupled sleep oscillations in REM-related processing is likely limited, since only (weak) coupling between delta/theta and beta/gamma components emerged during this sleep stage.

While variability is typically treated as a nuisance or noise factor in neuroscience, proper understanding of a phenomenon requires scrutiny of both the group mean and its natural variation (Kanai and Rees, 2011; Seghier and Price, 2018). (This is clearly illustrated by the HPC maps of Fig. 5A, where group means do not describe any individual particularly well.) More specifically, the degree of explanatory power that can be ascribed to a coupling phenomenon occurring in either 25 or 75% of individuals is decidedly different, even though group statistics might be similar. Analogously, observing a coupling phenomenon in either 1 or 9 out of 10 recording sites places different constraints on the functional role this coupling may play. While these additional layers of complexity present novel challenges, these obstacles can only be overcome by being aware of their existence in the first place.

## 5 Conclusions

While hippocampal sleep oscillations are thought to play a central role in systems memory consolidation, little is known about how coordinated rhythmic brain activity is organized in human HPC. Our findings reveal that such oscillatory interactions occur in remarkably diverse ways. While the causes of this heterogeneity remain to be determined, we believe that characterizing both the commonalities and variability of the sleep oscillatory landscape will contribute to a deeper understanding of sleep’s role in memory consolidation, and the function of sleep more broadly.

## Acknowledgements

This work was supported by the German Research Foundation (FE366/9-1 to J.F.) and a Wellcome Trust/Royal Society Sir Henry Dale Fellowship (107672/Z/15/Z to B.P.S.). The funders had no role in study design, data collection and analysis, decision to publish, or preparation of the manuscript.

