## Supplementary Materials for "Heterogeneous profiles of coupled sleep oscillations in human hippocampus"

### Heterogeneous profiles of coupled sleep oscillations in human hippocampus – Supplementary Information

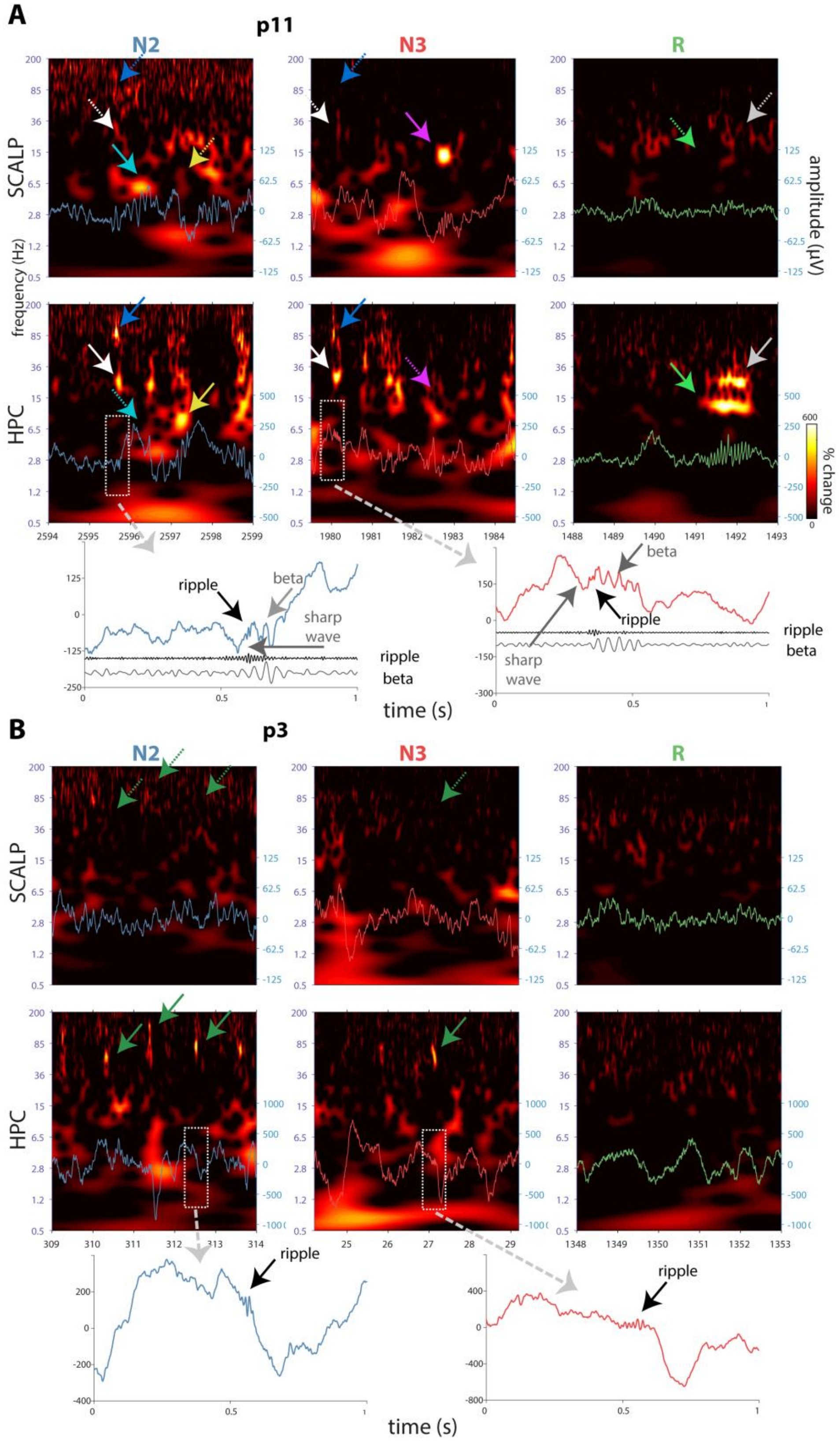

**Supplementary Figure 1 (previous page). Additional examples of scalp and hippocampal sleep activity.** Shown are 5 s segments of artifact-free data from two patients. Solid and dashed arrows indicate brain site with stronger and weaker effects, respectively. (A) Patient p11 (also shown in Fig. 4BC) expressed bouts of N2 scalp theta, without corresponding theta activity in HPC (light blue), and vice versa (yellow). Similarly, NREM spindle activity could be seen at the scalp but not in HPC (magenta), while clear HPC spindles were sometimes seen during REM (light green). Oscillatory beta (white) and ripple (dark blue) activity were seen in HPC during both N2 and N3, as further illustrated in the insets (beta bandpass: 20-50 Hz; ripple bandpass: 70-200 Hz). These ripples appeared on top of slower waveforms resembling sharp waves. Beta activity sometimes coincided with spindle activity (gray), likely reflecting a spindle harmonic due to nonsinusoidal waveform shape. HPC contact from anterior HPC. (B) Patient p3 showed visually apparent bursts of HPC ripple activity (dark green) of an oscillatory nature (insets). Also noteworthy is that N3 SO activity appears more prominent in HPC than at the scalp, and that SO-like activity can be seen in the HPC during REM. HPC contact from anterior HPC.

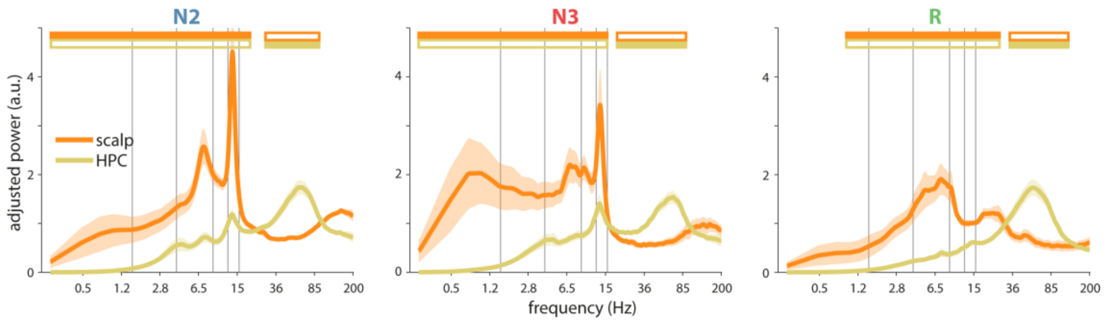


**Supplementary Figure 2. Comparison of scalp and hippocampal spectra.** Error shading: standard error of the mean across patients. Horizontal color bars above plots indicate significant (P<0.05) regional differences (cluster-based permutation test). Filled color reflects region with greater power. Note that while scalp and HPC signals have different amplitudes and are recorded using electrodes with different properties, the slope removal approach facilitates direct comparison between these sites.

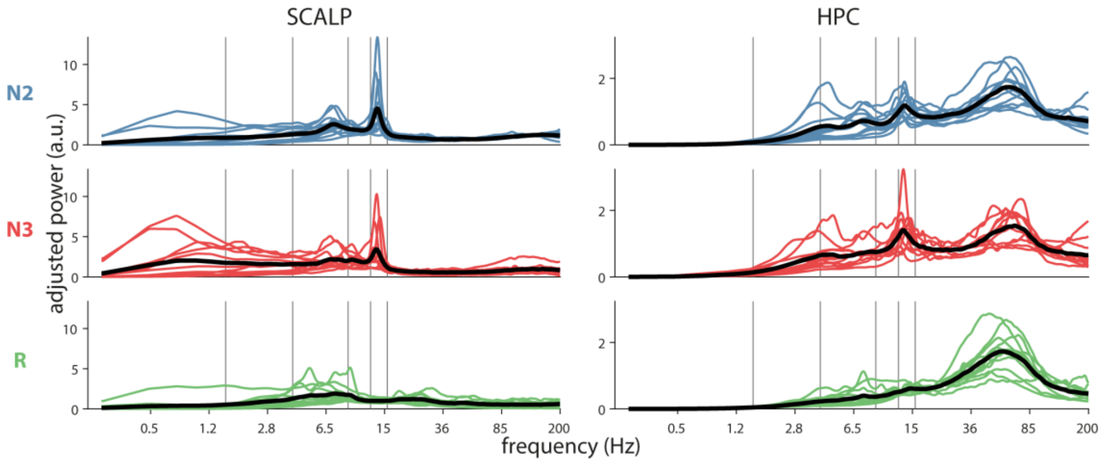


**Supplementary Figure 3. Individual scalp and hippocampal spectra of all patients.** Colored lines indicate single-patient scalp (left) and HPC spectra averaged across available channels (right). Black line indicates group mean.

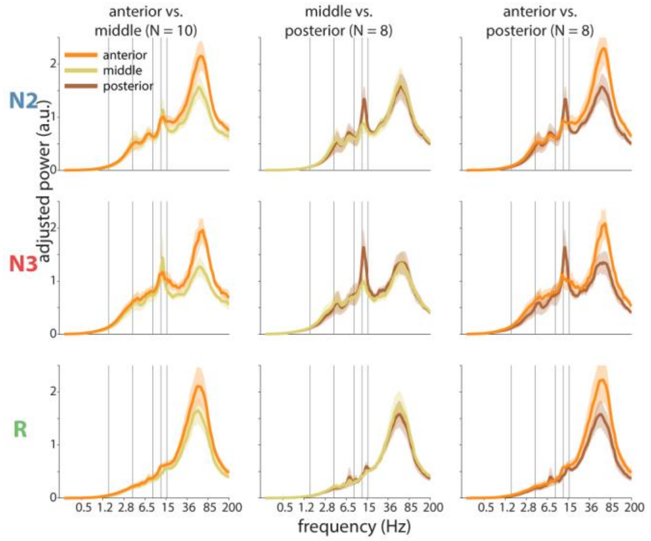


**Supplementary Figure 4. Comparison of anterior, middle, and posterior hippocampal spectra.** Single-patient HPC spectra averaged across available channels per region, followed by averaging across patients. Note the different sample sizes for different comparisons due to variable electrode coverage. Error shading: standard error of the mean across patients. No significant regional differences emerged (cluster-based permutation test).

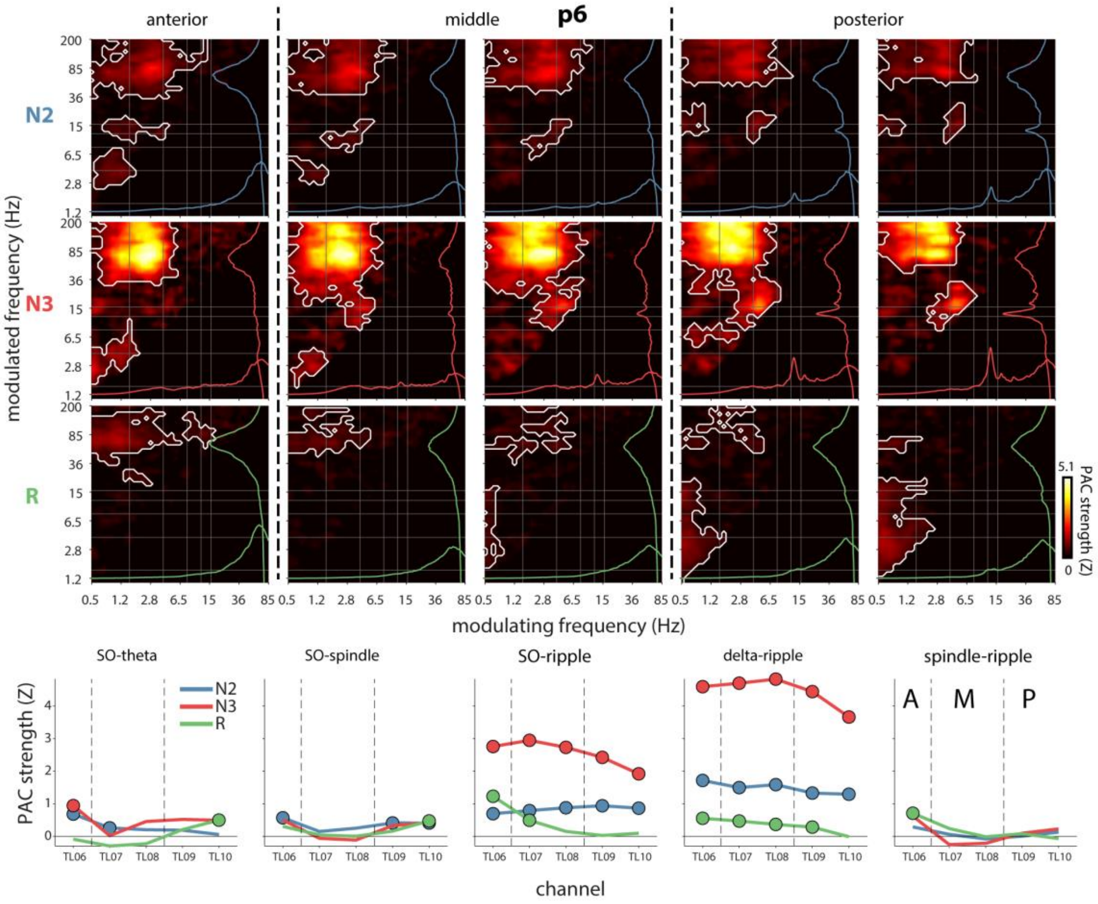


**Supplementary Figure 5.** Phase-amplitude coupling along hippocampal axis for additional patient. Top: Coupling strengths for all HPC contacts. White outlines indicate clusters of significantly higher than zero coupling (P<0.05, cluster-based permutation test). Bottom: Coupling strengths for five frequency pairs. Presence of marker indicates frequency pair is part of significant cluster in top panel. Dashed lines separate anterior (A), middle (M), and posterior (P) channels.

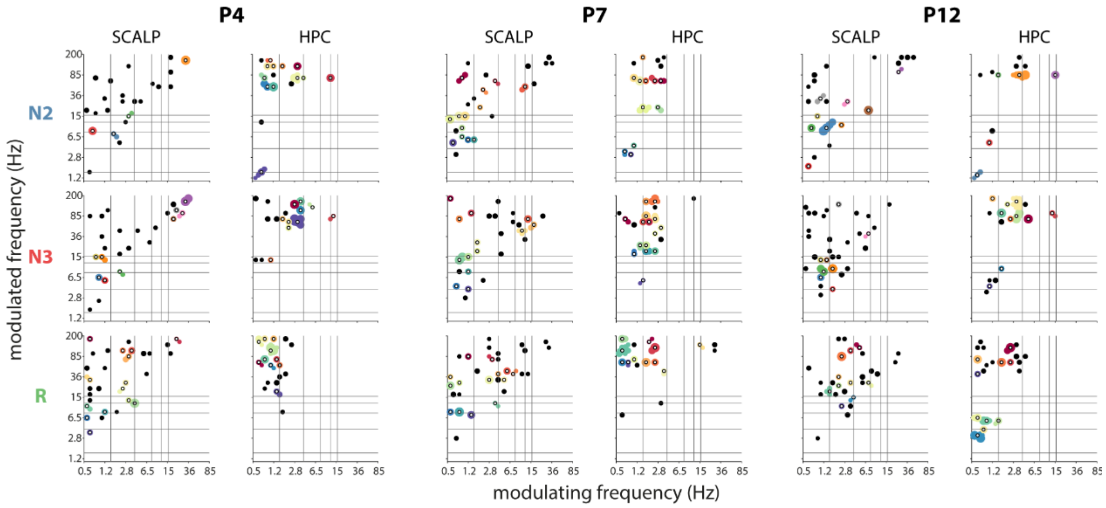


**Supplementary Figure 6. Examples of density-based clustering.** Every dot indicates a suprathreshold (adaptive approach) local maximum in coupling strength submitted to the DBSCAN algorithm. Colored dots belong to a cluster, with different colors for different clusters. (Color coding different for each panel.) Small open dots indicate cluster centers. Black dots indicate points not assigned to cluster.

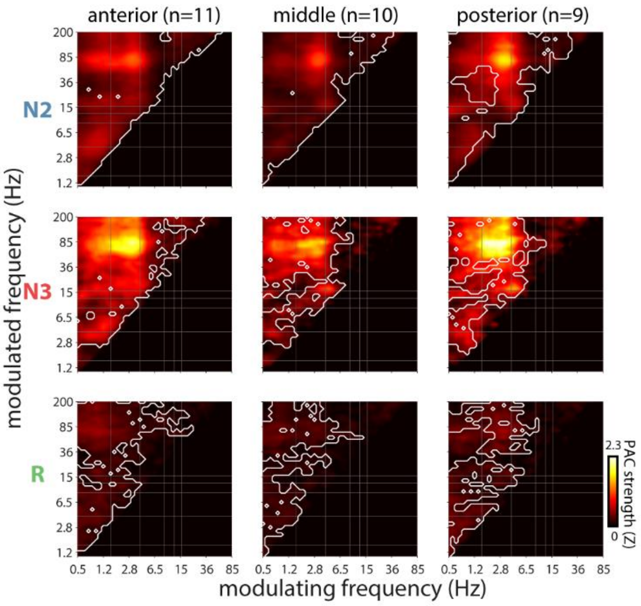


**Supplementary Figure 7. Group-level phase-amplitude coupling for hippocampal regions.** Mean coupling strength across patients for anterior, middle, and posterior HPC. White outlines indicate clusters of significantly higher than chance coupling (P<0.05, cluster-based permutation test). Note different sample sizes for each region due to variable electrode coverage.

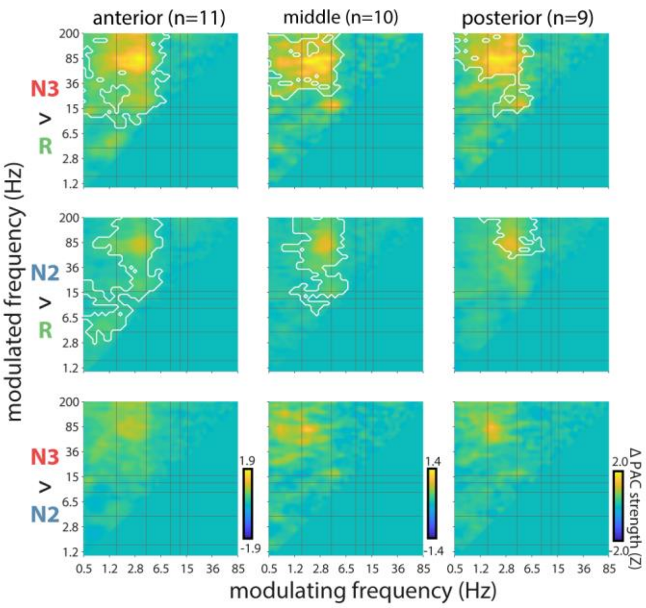


**Supplementary Figure 8. Stage differences in phase-amplitude coupling for hippocampal regions.** Mean difference in coupling strength between sleep stages for anterior, middle, and posterior HPC. White outlines indicate significant positive clusters (P<0.05, cluster-based permutation test; no negative clusters present). Note different sample sizes for each region due to variable electrode coverage.

| | | MNI | | |
|---|---|---|---|---|
| patient/ channel | region | x | y | z |
| p1 | RIGHT | | | |
| TR07 | P/wm | 32 | -31 | -3 |
| p2 | LEFT | | | |
| TL05 | A | -24 | -12 | -26 |
| TL06 | A | -25 | -16 | -22 |
| TL07 | A | -24 | -19 | -18 |
| TL08 | M | -24 | -22 | -16 |
| TL09 | M | -24 | -26 | -11 |
| p3 | RIGHT | | | |
| TRb02 | A | 27 | -9 | -27 |
| TRb03 | A | 29 | -11 | -20 |
| TRb04 | A | 29 | -16 | -17 |
| TRb05 | M | 28 | -20 | -13 |
| TRb06 | M/wm | 30 | -25 | -9 |
| TRb07 | P/wm | 30 | -29 | -5 |
| p4 | LEFT | | | |
| TL02 | A | -26 | -14 | -21 |
| TL03 | A | -26 | -17 | -18 |
| TL04 | M | -26 | -20 | -16 |
| TL05 | M | -26 | -24 | -13 |
| TL06 | P/wm | -27 | -28 | -10 |
| TL07 | P/wm | -28 | -32 | -5 |
| TL08 | P/wm | -29 | -35 | -1 |
| p5 | RIGHT | | | |
| TR01 | A | 26 | -12 | -24 |
| TR02 | A | 27 | -15 | -21 |
| TR03 | A | 27 | -18 | -18 |
| TR04 | A | 27 | -22 | -14 |
| TR05 | M | 27 | -24 | -11 |
| TR06 | M | 27 | -29 | -9 |
| TR07 | M | 27 | -33 | -4 |
| TR08 | P | 29 | -35 | 2 |
| p6 | LEFT | | | |
| TL06 | A | -26 | -17 | -19 |
| TL07 | M | -27 | -21 | -16 |
| TL08 | M | -27 | -25 | -14 |
| TL09 | P | -28 | -28 | -12 |
| TL10 | P | -29 | -24 | -8 |
| p7 | RIGHT | | | |
| TR04 | A | 22 | -16 | -22 |
| TR05 | A | 22 | -20 | -21 |
| TR06 | M | 22 | -24 | -19 |
| TR07 | M | 24 | -27 | -14 |
| TR08 | M | 25 | -31 | -9 |
| TR09 | P | 28 | -35 | -5 |
| TR10 | P | 29 | -39 | -2 |
| p8 | RIGHT | | | |
| TR03 | A | 28 | -8 | -21 |
| TR04 | A | 28 | -11 | -19 |
| TR05 | M/lv | 28 | -15 | -17 |
| TR06 | M/lv | 29 | -19 | -15 |
| TR07 | M/wm | 30 | -23 | -12 |
| TR08 | P/wm | 31 | -29 | -8 |
| p9* | LEFT | | | |
| TLa01 | A | -27 | -13 | -25 |
| TLa02 | A | -31 | -15 | -24 |
| p10 | LEFT | | | |
| TL02 | A/lv | -23 | -13 | -17 |
| TL03 | M/lv | -23 | -15 | -14 |
| p11 | LEFT | | | |
| TL03 | A | -33 | -19 | -17 |
| TL04 | M | -33 | -23 | -16 |
| TL05 | M | -32 | -26 | -14 |
| TL06 | P | -32 | -30 | -11 |
| TL07 | P | -33 | -34 | -8 |
| p12 | RIGHT | | | |
| TR04 | A | 23 | -5 | -40 |
| TR05 | A | 26 | -17 | -34 |
| TR06 | M | 27 | -22 | -30 |
| TR07 | M | 27 | -25 | -25 |
| TR08 | P | 27 | -29 | -19 |
| TR09 | P | 26 | -31 | -14 |
| TR10 | P | 28 | -35 | -9 |

**Supplementary Table 1. Locations of included hippocampal contacts.** Electrodes were included when located wholly within HPC gray matter, on the HPC gray/white matter (wm) border, or the HPC gray matter/lateral ventricle (lv) border. Contacts were classified as anterior (A), middle (M) or posterior (P) based on individual anatomy. MNI coordinates after warping to MNI template are also provided. * indicates patient with lateral implantation.

| | **N2 scalp** | | | **N2 HPC** | | | **N3 scalp** | | | **N3 HPC** | | |
|---|---|---|---|---|---|---|---|---|---|---|---|---|
| frequency pair | f1 | f2 | N | f1 | f2 | N | f1 | f2 | N | f1 | f2 | N |
| SO–delta | | | | 1.0–1.2 | 4.5–5.1 | 4 | 0.6–0.7 | 4.0 | 4 | | | |
| | | | | | | | 1.0 | 3.5–4.5 | 3 | | | |
| | | | | | | | 1.3–1.7 | 3.5 | 3 | | | |
| SO–theta | 0.6 | 4.5–5.8 | 4 | | | | 1.0 | 7.4 | 5 | | | |
| | 1.3 | 5.8 | 4 | | | | 1.3 | 7.4–8.3 | 5 | | | |
| SO–theta/slow spindle | 0.8 | 8.3–9.4 | 4 | | | | | | | | | |
| | 0.6 | 8.3–9.4 | 3 | | | | | | | | | |
| | 1.5 | 8.3–10.6 | 3 | | | | | | | | | |
| SO–fast spindle | 1.0 | 13.6–15.3 | 5 | | | | **1.0** | **15.3** | **9** | | | |
| SO–beta/gamma | 0.9–1.2 | 36.1 | 3 | 0.9 | 75.2–85.0 | 4 | | | | 1.3 | 28.2–40.8 | 4 |
| | | | | 0.9–1.0 | 156.7–177.0 | 3 | | | | 0.8 | 75.2 | 4 |
| | | | | | | | | | | 0.9–1.2 | 19.6 | 3 |
| | | | | | | | | | | 1.3–1.5 | 156.6 | 3 |
| delta–spindle/beta | | | | 1.5–1.7 | 19.6 | 3 | 1.9 | 17.3 | 3 | 2.8–3.1 | 19.6–25.0 | 5 |
| | | | | | | | | | | 1.7 | 28.2–36.1 | 4 |
| delta–beta/gamma | 1.9 | 40.8 | 3 | **3.1** | **75.2** | **12** | | | | **2.8** | **75.2** | **11** |
| | | | | **3.1** | **46.1** | **7** | | | | **1.5** | **85.0** | **8** |
| | | | | 1.5 | 75.2 | 5 | | | | **2.5** | **46.1** | **7** |
| | | | | 2.8–3.1 | 108.5 | 5 | | | | 2.2–2.8 | 36.1–46.1 | 6 |
| | | | | 2.8–3.1 | 138.6 | 5 | | | | 3.1–3.6 | 138.6 | 6 |
| | | | | | | | | | | 2.5 | 122.6–138.6 | 5 |
| | | | | | | | | | | 4.5 | 75.2 | 4 |
| delta/theta–spindle | | | | 4.0–5.1 | 15.3–19.6 | 4 | | | | 4.5 | 15.3–19.6 | 5 |
| theta–gamma | 8.3 | 58.9–66.5 | 3 | | | | | | | | | |
| slow spindle/alpha–gamma | | | | | | | 10.6–12.0 | 138.6 | 3 | | | |
| spindle–ripple | | | | 13.6 | 75.2–85.0 | 3 | | | | 12.1 | 85.0 | 3 |
| beta–gamma | **36.1** | **156.6–177.0** | **7** | | | | **22.1** | **108.5** | **7** | | | |
| | 22.1–25.0 | 108.5 | 6 | | | | **32.0** | **156.6** | **7** | | | |
| | 19.6 | 122.6 | 6 | | | | 17.3–19.6 | 66.5 | 4 | | | |
| | 22.1 | 138.6 | 6 | | | | | | | | | |

**Supplementary Table 2. Prevalence of NREM coupling following density-based clustering (adaptive approach).** Indicated is how many individuals (N) expressed coupling for each combination of a modulating frequency range (f1) with a modulated frequency range (f2). Only effects present in at least 3 individuals (25%) are shown. Effects present in more than half of the sample (>6) are indicated in bold.
